# Evolution of phenotypic variance provides insights into the genetic basis of adaptation

**DOI:** 10.1101/2021.01.19.427260

**Authors:** Wei-Yun Lai, Viola Nolte, Ana Marija Jakšić, Christian Schlötterer

**Affiliations:** Institut für Populationsgenetik, Vetmeduni Vienna, Vienna, Austria; Vienna Graduate School of Population Genetics, Vetmeduni Vienna, Vienna, Austria

**Keywords:** Phenotypic variance, temperature adaptation, *Drosophila simulans*, experimental evolution

## Abstract

Most traits are polygenic and the contributing loci can be identified by GWAS. Their adaptive architecture is, however, difficult to characterize. Here, we propose to study the adaptive architecture of traits by monitoring the evolution of their phenotypic variance during adaptation to a new environment in well-defined laboratory conditions. Extensive computer simulations show that the evolution of phenotypic variance in a replicated experimental evolution setting can distinguish between oligogenic and polygenic adaptive architectures. We compared gene expression variance in male *Drosophila simulans* before and after 100 generations of adaptation to a novel hot environment. The variance change in gene expression was indistinguishable for genes with and without a significant change in mean expression after 100 generations of evolution. We suggest that a majority of adaptive gene expression evolution can be explained by a polygenic architecture. We propose that tracking the evolution of phenotypic variance across generations can provide an approach to characterize the adaptive architecture.

**Significant Statement:** It is widely accepted that most complex traits have a polygenic basis. Nevertheless, it is difficult to predict which of these loci are responding to selection when a population is exposed to a new selection regime. To address this situation, we propose to infer the adaptive architecture for traits by tracking the evolution of their phenotypic variance during adaptation to a new environment. As a case study, we analyze the evolution of gene expression variance in outbred *Drosophila simulans* populations adapting to a new temperature regime to infer the genetic architecture of adaptive gene expression evolution. We suggested that the adaptive gene expression evolution is better explained by a polygenic architecture.

## Introduction

It is widely accepted that most complex traits have a polygenic basis (Ayroles *et al*., 2009; Boyle *et al*., 2017; Liu *et al*., 2019). Nevertheless, it is difficult to predict which of these loci are responding to selection when a population is exposed to a new selection regime. If pleiotropic constraints are strong, only a small subset of the contributing genes are free to respond to selection. Hence, the genetic basis of the adaptive response of a complex trait (i.e. adaptive architecture (Barghi *et al*., 2020)) may differ substantially from the genetic architecture. Characterizing the adaptive architecture by mapping selected loci is not easy, in particular when more than a handful of genes are involved. To circumvent this problem, we introduce an approach, analogous to the Castle-Wright estimator (Castle, 1921), to infer the complexity of the adaptive architecture (i.e. simple with few contributing loci or complex with a polygenic basis). We propose to study the evolution of phenotypic variance, which may provide some insights into the key parameters of the adaptive architecture.

The phenotypic variance of a quantitative trait is a key determinant for its response to selection. It can be decomposed into genetic and environmental components (Falconer & Mackay, 1963). Over the past years, mathematical models have been developed to describe the expected genetic variance of a quantitative trait under selection and its maintenance in evolving populations (Bulmer, 1972; Chevalet, 1994; Kimura and Crow, 1964; Turelli, 1984). For infinitely large populations and traits controlled by many independent loci with infinitesimal effect, changes in trait optimum are not expected to affect the phenotypic variance (Lande, 1976). A much more complex picture is expected when the effect sizes are not equal, the population size is finite, or the traits have a simpler genetic basis (Barton & Turelli, 1987; Keightley & Hill, 1989; Barton & Keightley, 2002; Jain & Stephan, 2015; Franssen *et al*., 2017; Hayward & Sella, 2019). For instance, for traits with oligogenic architecture, the genetic variance could drop dramatically during adaptation, while with polygenic architectures, only minor effects on the variance are expected (Barton et al., 2017; Franssen et al., 2017; Jain and Stephan, 2015). These insights suggest that a time-resolved analysis of phenotypic variance has the potential to shed light onto the complexity of the underlying adaptive architecture.

Despite its potential importance for the understanding of adaptation, we are faced with the situation that few empirical data are available for the evolution of phenotypic variance. The use of natural populations to study changes in phenotypes, and even more so phenotypic variances, is limited as the environmental heterogeneity cannot be controlled and common garden experiments are required to study the phenotypic variance, which is not feasible for many species. A complementary approach to study the evolution of phenotypic variance is experimental evolution (Kawecki *et al*., 2012). With replicated populations starting from the same founders and evolving under tightly controlled environmental conditions, experimental evolution provides the opportunity to study the evolution of phenotypic variance.

Most experimental evolution studies in sexual populations focused on the evolution of phenotypic means, rather than variance (e.g.: Chippindale *et al*., 1996; Burke *et al*., 2010; Mallard *et al*., 2018; Jakšić *et al*., 2020). A notable exception is a study which applied fluctuating, stabilizing and disruptive selection to a small number of wing shape-related traits (Pélabon *et al*., 2010). Other studies tracked the variance evolution of behavioral and cranial traits under directional selection in mice (Careau *et al*., 2015; Penna *et al*., 2017). Instead of looking at a preselected subset of phenotypes which limits the generality, we will focus on gene expression, a set of molecular phenotypes, which can be easily quantified as microarrays and, more recently, RNA-Seq have become available. Importantly, the expression levels of genes exhibit the same properties (e.g.: continuality and normality) as other complex quantitative traits (Mackay *et al*., 2009). Thus, gene expression has also been widely employed to study the adaptation of locally adapted populations (e.g.: Romero *et al*. 2012; Sork 2017; Signor and Nuzhdin 2018) or ancestral and evolved populations in the context of experimental evolution (e.g.: Lenski *et al*. 1994; Ferea *et al*. 1999; Huang and Agrawal 2016; Mallard *et al*. 2018).

In this study, we performed forward simulations that match not only essential design features of typical experimental evolution studies, but also incorporate realistic parameters of the genetic architecture. We recapitulate the classic results that even a moderately polygenic architecture is associated with a high stability of the phenotypic variance of selected traits across different phases of adaptation and evaluate the predictive performance under different scenarios. Applying this insight to a recently published dataset (Lai & Schlötterer, 2022), we show that for putatively selected genes (DE genes), their average changes in expression variance were indistinguishable from genes without changes in mean expression. We propose that this pattern reflects a polygenic basis of adaptive gene expression evolution.

## Results

The central idea of this study is that the complexity of adaptive traits can be inferred from the trajectory of the phenotypic variance during adaptation: the phenotypic variance remains relative stable for a trait with polygenic (infinitesimal) architecture while it changes across generations for a trait with oligogenic architecture. Although this prediction has been illustrated in multiple theoretical and simulation studies (e.g.: (Jain & Stephan, 2015; Barton *et al*., 2017; Franssen *et al*., 2017)), its application to empirical data has been limited. A recent experimental evolution study reporting the phenotypic variance of more than 10,000 gene expression traits before and after 100 generations of adaptation to a novel environment (Lai & Schlötterer, 2022) provides an opportunity to fill this gap. Since gene expression can be viewed as quantitative traits (Ayroles *et al*., 2009; Hill *et al*., 2021), we apply the theoretical predictions to the empirical data to infer the genetic basis of adaptive gene expression evolution. Hence, as the first step of this study, we explored to what extent theoretical predictions can be generalized to expression traits obtained from a typical experimental evolution setting by considering a broad parameter space and accounting for linkage. Using population genetic parameters (e.g. number of starting haplotypes, effective population size) estimated from the above-mentioned evolution experiment, we simulated 1000 neutral and 1000 selected expression traits. To account for experimental and environmental noise, we used an empirical distribution of heritabilities (estimated from gene expression measurements across F1 families; see supplementary information). The simulated selection regime consists of a mild/distant shift in trait optimum with weak/intermediate/strong stabilizing selection (Figure 1a and Supplementary Figure S4) (Hayward & Sella, 2019). We assumed additivity and a negative correlation between the ancestral allele frequency and the effect size of contributing loci (Otte et al. 2020) (Figure 1b). With three different distributions of effect size (Figure 1c), we investigated how the number of contributing loci affects the evolution of gene expression variance with and without selection.

**Figure 1.**
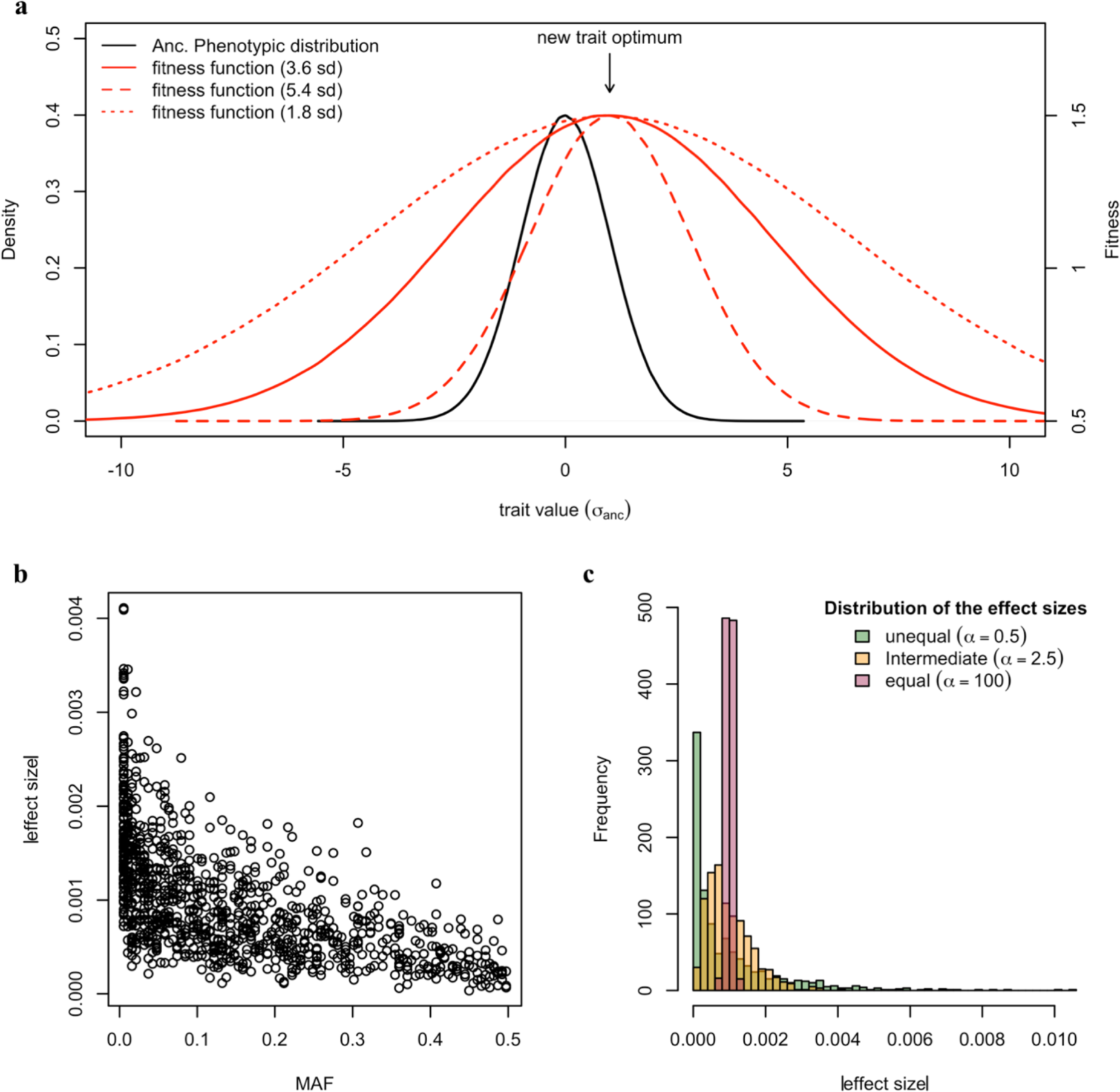
Simulating polygenic adaptation to a shift in trait optimum with different parameter combinations. **a.** For the computer simulations we consider a quantitative trait (in black) experiencing a sudden shift in trait optimum under stabilizing selection. The underlying fitness functions are illustrated in red. The new trait optimum is shifted from the ancestral trait mean by one/three standard deviation of the ancestral trait distribution. The strength of stabilizing selection is modified by changing the variance of the fitness function: 1.8, 3.6 and 5.4 standard deviations of the ancestral trait distribution. **b.** The negative correlation between the allele frequencies and the effect sizes (r = -0.7, estimated in Otte *et al*. (2020)). We consider such negative correlation when assigning the effect sizes to variants underlying a simulated trait. **c.** The distribution of effect sizes of the contributing loci is determined by the shape parameter (α) of gamma sampling process (α = 0.5, 2.5 and 100).

We monitored the change in phenotypic variance over 100 generations, which was sufficient to reach the trait optimum for most parameter combinations (Supplementary Figure S5). We compared the change in variance relative to the start of the experiment in populations with and without selection. First, we studied a mild (one standard deviation of the ancestral phenotypic distribution) shift in trait optimum. As expected for a founder population derived from a substantially larger natural population, we find that even under neutrality, the phenotypic variance does not remain constant, but gradually decreases during 100 generations of experimental evolution (Figure 2). This pattern is unaffected by the genetic architecture of the neutral traits. We explain this loss of variance by the fixation of variants segregating in the founder population and the fact that we did not simulate new mutations, as they do not contribute to adaptation in such short time scales (Burke *et al*., 2010). Although our simulations used moderate population sizes, they nicely recapitulate the patterns described for populations without drift (Jain & Stephan, 2015; Barton *et al*., 2017). A pronounced drop in phenotypic variance is observed when a trait is approaching a new optimum with few contributing loci (Figure 2). When more loci (with smaller effects) are contributing to the selected phenotype, the difference to neutrality becomes very small (Figure 2). In addition to the number of contributing loci, the heterogeneity in effect size among loci and the shape of the fitness function have a major impact. The larger the difference in effect size is, the more pronounced was the influence of the number of contributing loci (Figure 2). The opposite effect was seen for the width of the fitness function – a wider fitness function decreased the influence of the number of contributing loci (Figure 2). Importantly, these patterns were not affected by the duration of the experiment - qualitatively identical patterns were seen at different time points until generation 200.

**Figure 2.**
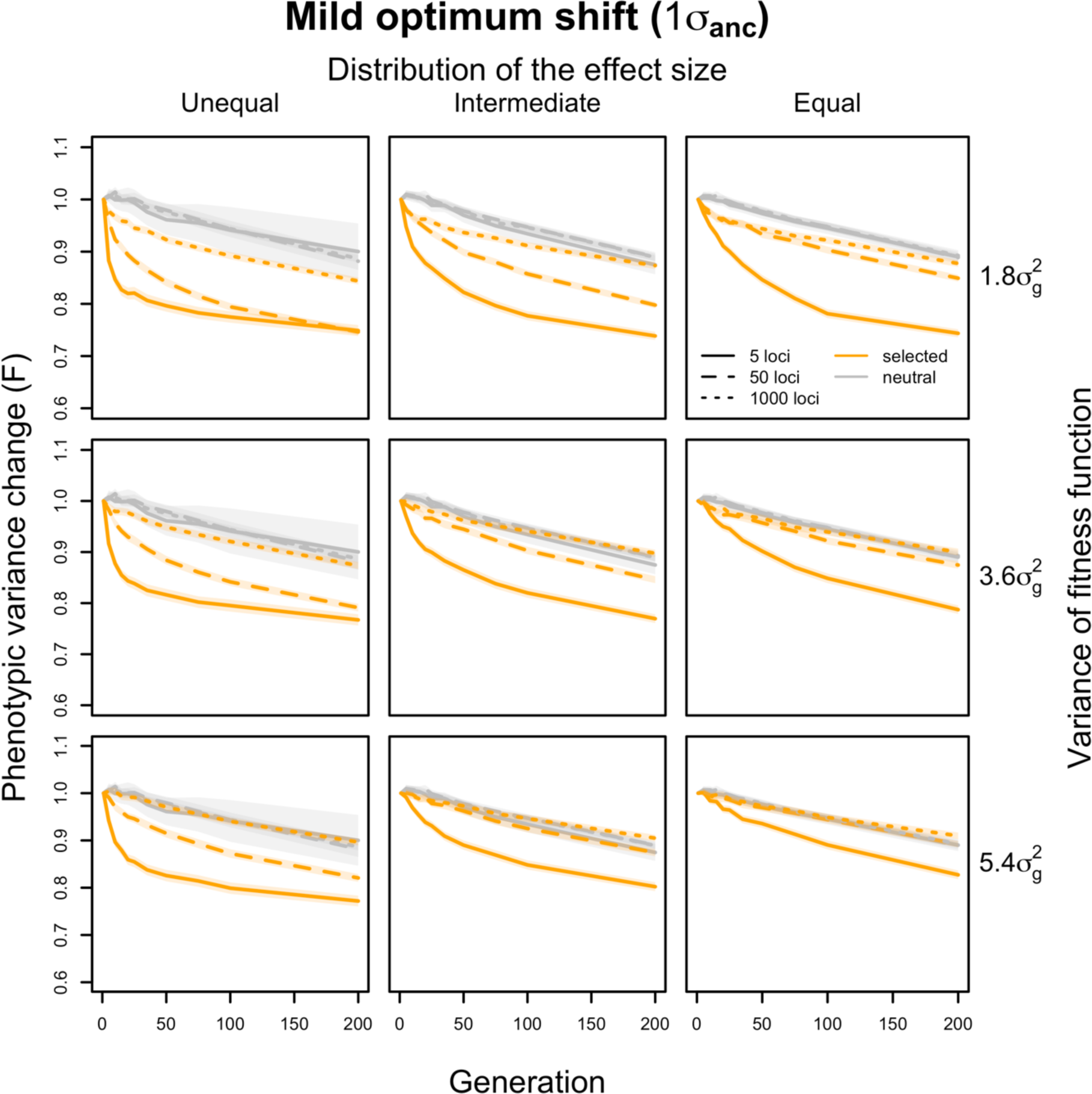
The trajectory of changes in phenotypic variance during adaptation to a mild optimum shift. The changes in phenotypic variance within 200 generations adapting to a moderate optimum shift (orange) are compared to the changes under neutrality (grey) on y axis. The average variance changes (F) of 1000 simulated traits is calculated as the ratio of phenotypic variance between each evolved time point (generation *x*) and the ancestral state (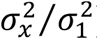). The translucent band indicates the 95% confidence interval for 1000 simulated traits. The simulations cover traits controlled by varying numbers of loci underlying the adaptation with three different distributions of effect sizes (columns) under different strength of stabilizing selection (rows). Only traits with the most (dotted lines, 1000 loci), intermediate (dash lines, 50 loci) and the least (solid lines, 5 loci) polygenic architectures are shown. In all scenarios, the variance of the trait decreases drastically when the adaptation is controlled by a small number of loci (orange solid lines; 5 loci). While, for traits with extremely polygenic basis, the phenotypic variance stays stable over time (orange dotted lines).

For a more distant trait optimum (three standard deviations of the ancestral phenotypic distribution), we noticed some interesting dynamics that were not apparent for a closer trait optimum (Supplementary Figure S6). The most striking one was the temporal heterogeneity of the phenotypic variance when few loci of unequal effects are contributing. During the early stage of adaptation, the variance increased and dropped later below the variance in the founder population. With an increasing number of contributing loci, this pattern disappeared and closely matched the neutral case (Supplementary Figure S6).

Overall, our simulations indicate that with a larger number of contributing loci the variance fitted the neutral pattern better. Modifying dominance did not change the overall patterns (Supplementary Figure S7). The large influence of key parameters of the adaptive architecture, in particular the number of contributing loci and their effect sizes, on the temporal phenotypic variance dynamics suggests that it should be possible to exploit this relationship to characterize the genetic basis of phenotypic adaptation.

A potentially interesting application of the relationship between number of contributing loci and evolution of variance is the inference of the adaptive architecture of a given trait. For a single selected trait, a F test contrasting the reduction of variance after adaptation against neutral loss of variance (∼0.9 after 100 generations in our simulations) can be used to infer the number of loci contributing to adaptive evolution. The lower bound for the number of contributing loci can be determined by the largest number of loci that results in a significant difference between a selected trait and the neutral expectation in all parameter combinations. Nevertheless, the power of this approach depends strongly on the sample size. When the entire population (N = 300) is phenotyped for the focal trait, the power to rule out an adaptive architecture of no more than 5 loci by significant decrease in variance of a selected trait is only 44%. This indicates the limited ability to distinguish polygenic from oligogenic architectures of single traits. With a more realistic sample size of 20 individuals the distinction is even more difficult such that we conclude that it is not possible to infer the adaptive architecture for a single trait when only two time points are available (Supplementary Figure S8).

Alternatively, it is possible to study multiple, presumably independent, selected phenotypes together. Assuming a similar genetic architecture of the selected traits, we showed that the distribution of variance changes of 1000 selected traits with a simple genetic architecture (five loci) can be distinguished from neutral changes. Even with a sample size of 20 individuals, as in our experimental data (see Materials and Methods), significant differences from neutral expectations can be detected for some parameters with a power close to 100% (Figure 3 and Supplementary Figure S9). This suggests that oligogenic and polygenic architectures of a group of selected traits can be distinguished experimentally even with moderate sample sizes.

**Figure 3.**
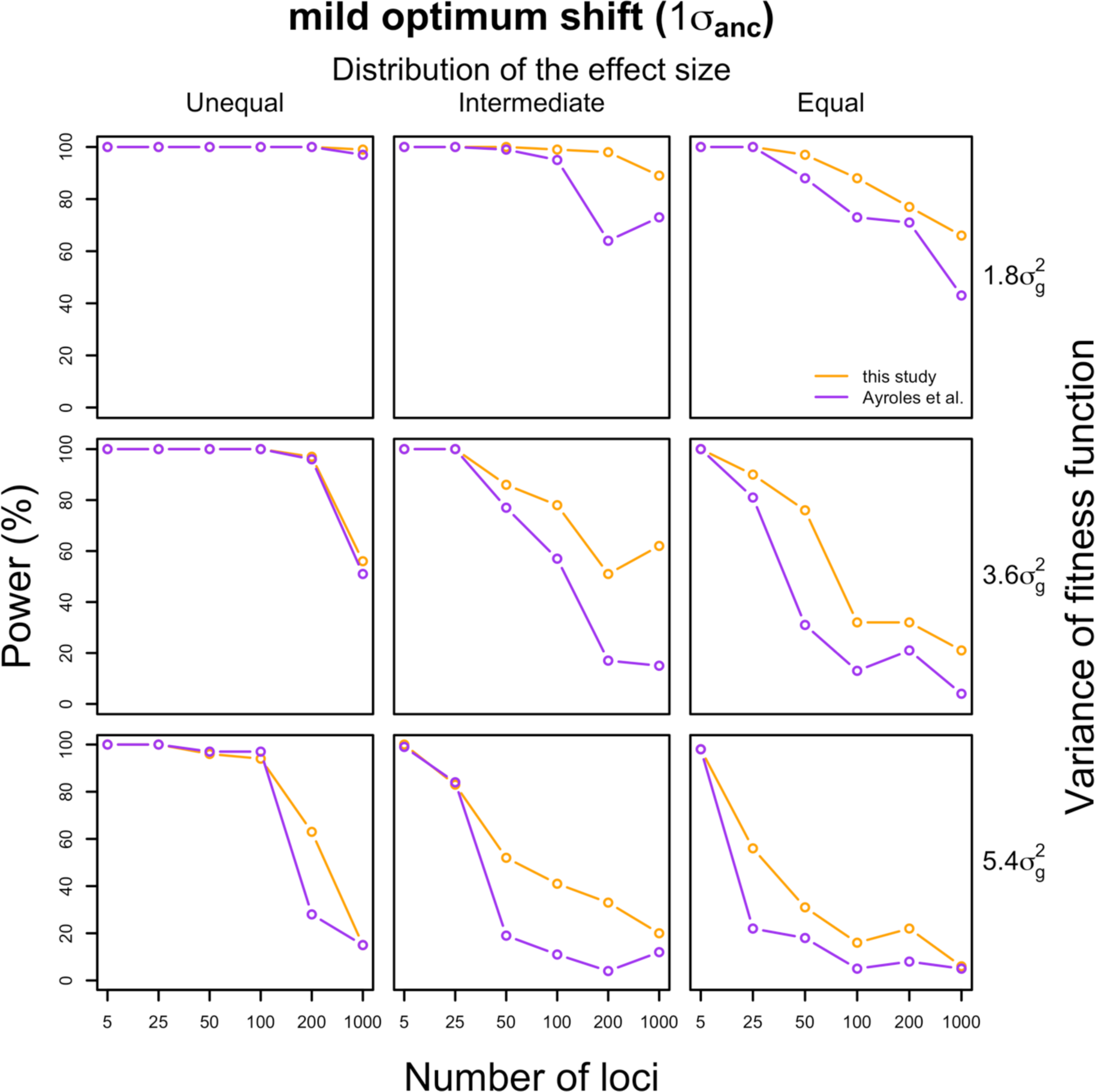
Power for detecting significant variance change between groups of selected and neutral traits. For each set of the 1000 traits controlled by different numbers of loci (x-axis) with varying effect sizes (columns) under each selection strength (rows), we calculated how often no difference in variance change is detected between 1000 neutral and 1000 selected traits after 100 generations (y-axis). The same genetic architecture is assumed for all 1000 selected traits with a sample size of 20. For each parameter combination, we have almost 100% power when all traits were controlled by few loci (5 loci). It gradually increases with an increasing number of loci (red). A lack of significant difference indicates polygenic architecture of the adaptive traits. The results remain unaffected when we used heritability estimates from Ayroles et al., (2009) (purple).

In the simulations, we fixed most experimental parameters such as population sizes, generation and sample sizes to reflect the experimental design of typical experimental evolution studies. Nevertheless, for future experimental evolution studies it is important to understand how the experimental design affects the power. We performed additional simulations to explore the impact of the sample size, number of generations and population size. As expected, the power increases with larger sample size, allowing to discriminate architectures with a larger number of contributing loci (Figure 4a). Later generations are also more informative than earlier generations (i.e., higher power to reject a larger number of loci) (Figure 4b). Larger population sizes could also improve the determination of polygenicity, as we find an increase in power for a population size of 1200 compared to 300 (Figure 4c). In summary, larger and longer evolution experiments with larger sample size are superior. They provide a better discrimination of genetic architectures with a larger number of contributing loci-i.e. a better distinction between oligogenic and polygenic architectures.

**Figure 4.**
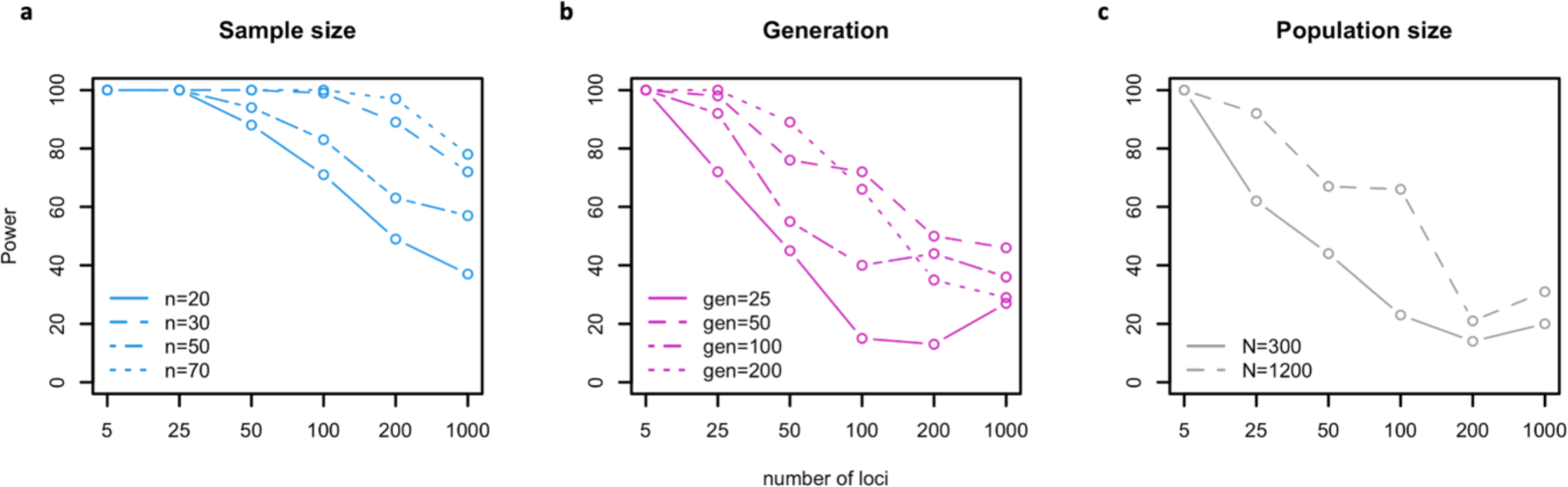
Power for detecting significant variance change between groups of selected and neutral traits under different a. sample sizes, b. generations and c. population sizes. For each set of the 1000 traits controlled by different numbers of loci (x-axis) under different experimental conditions, we calculated how often a difference in variance change is detected between 1000 neutral and 1000 selected traits (y-axis). The same genetic architecture is assumed for all 1000 selected traits. **a.** The power gradually increases with larger sample sizes. **b.** Given the same number of loci; a higher power is obtained when the experiment continues for more generations. **c.** Experiments with larger population size (N=1200) have a higher power than those with a smaller population size (N=300). Overall, larger and longer evolution experiments with more phenotyped samples provide more power to reject simpler adaptive architectures and thus provide more confidence in a highly polygenic architecture when we observed no significant difference in variance.

As a case study, we investigated the evolution of gene expression variance in replicated populations evolving in a new hot temperature regime and inferred the adaptive architecture of gene expression traits. The evolved populations were derived from the same ancestral population but evolved independently for more than 100 generations in a novel temperature regime with daily temperature fluctuations between 18 and 28°C (Figure 5a). Rather than relying on pooled samples which only allow mean estimates, we quantified gene expression of 19-22 individuals from reconstituted ancestral populations (F0) and two evolved populations (F103) in a common garden setup (Lai & Schlötterer, 2021). Principal Component Analysis (PCA) indicated that 11.9% of the variation in gene expression can be explained by the first PC which separates evolved and ancestral populations, reflecting clear adaptive gene expression changes in response to the novel hot temperature regime (Figure 5b). The means and variances of the expression of each gene were estimated and compared between the two reconstituted ancestral populations and the two evolved populations separately (see Materials and methods). Due to the usage of different lot numbers for the RNA-Seq library preparation (Table S1), we only contrasted ancestral and evolved samples generated with the same lot number (see Materials and methods) to avoid any unnecessary confounding effects.

**Figure 5.**
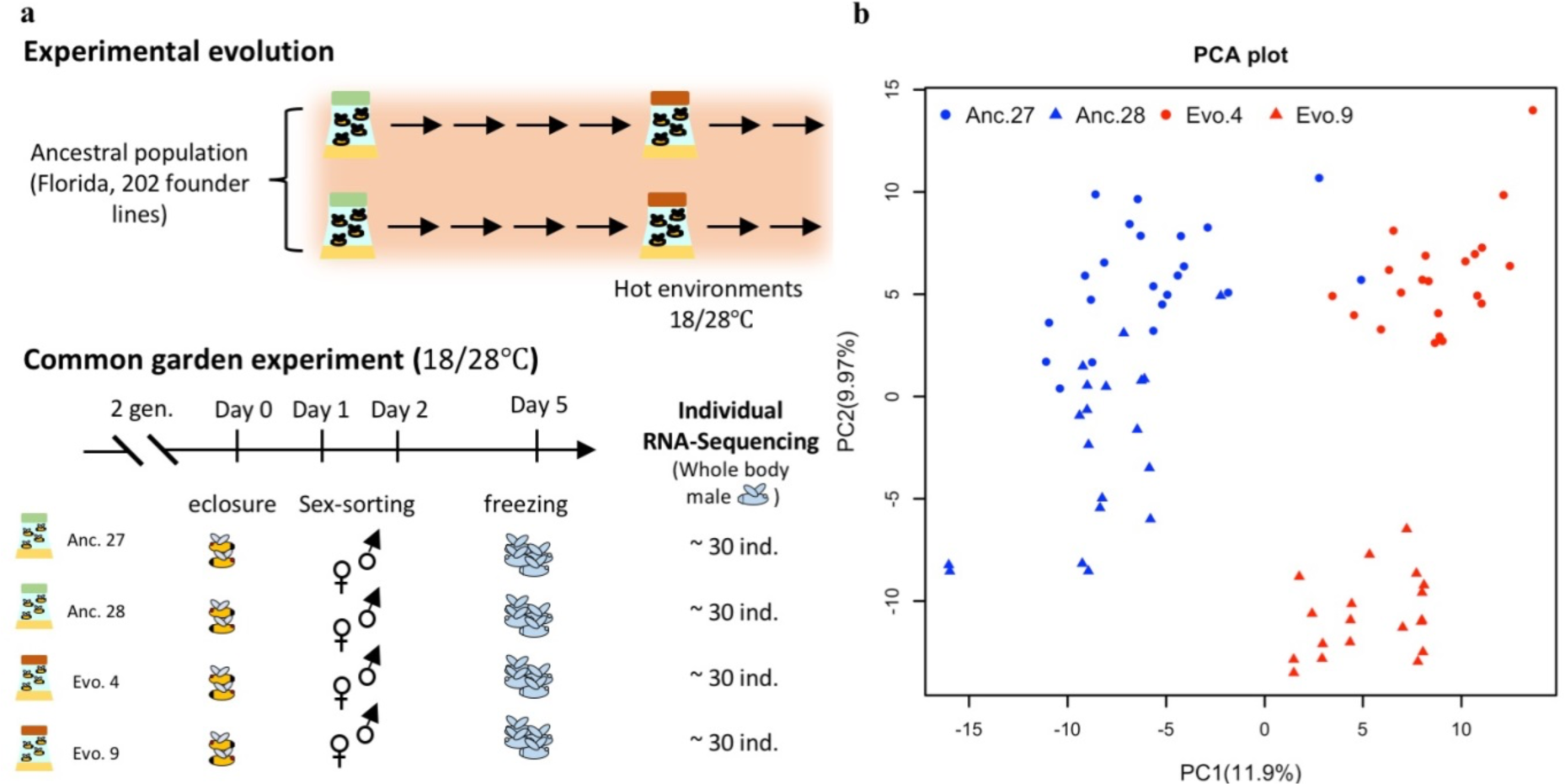
Schematic overview of the experimental procedures (a) and the divergence in gene expression during experimental evolution (b). **a.** Experimental evolution: starting from one common founder population, two replicate populations evolved for 100 generations in a hot laboratory environment fluctuating between 18 and 28°C. Common Garden Experiment: after 100 generations, the two evolved replicate populations were maintained together with the reconstituted ancestral population for two generations in a hot laboratory environment fluctuating between 18 and 28°C. After this common garden procedure, males from each population were subjected to RNA-Seq. **b.** Principal Component Analysis (PCA) of the transcriptomic profiles of individuals from the ancestral population (blue) and the hot-evolved population (red). Circles indicate individuals of the first replicate (Anc. No. 27 and Evo. No. 4). Triangles represent individuals of the second replicate (Anc. No. 28 and Evo. No. 9). The two replicates were made with two different batches of library cards for RNA-Seq library preparation. The impact of the library card batch can be seen from the separation of the ancestral replicates which were reconstituted from the same iso-female lines.

The comparison of ancestral and evolved populations identified 2,775 genes in the first population and 2,677 genes in the second population which significantly changed mean expression in the evolved flies (FDR<0.05, (Lai & Schlötterer, 2022)). 1,172 differentially expressed genes are shared between two populations (Figure 6a). The concordance of both populations and the similarity to previous studies of the same evolution experiment (Hsu *et al*., 2020; Jakšić *et al*., 2020; Lai & Schlötterer, 2022) suggests that most of the altered expression means are mainly driven by selection, rather than by drift. We scaled the gene expression change with the standard deviation in the ancestral population to approximate the selection strength on each gene. The differentially expressed genes in both populations showed a broad distribution of expression change, but the averaged mean expression changed by one standard deviation (Figure 6b), which is unlikely to be reached by drift based on our simulations (less than 1% of the neutral traits drifted by one standard deviation of the ancestral phenotypic variance). Assuming that these putative adaptive gene expression phenotypes reached the new trait optimum, this corresponds, on average, to a mild shift in trait optimum in our computer simulations. Reasoning that genes with significant mean expression changes are under selection and the rest of the transcriptome is not responding to the new environment (neutral), we compared the evolutionary changes in expression variance of genes with and without significant changes in expression mean (DE and non-DE genes), to characterize the adaptive architecture of gene expression. The underlying hypothesis is that a simple adaptive architecture will be associated with a significant drop in the expression variance of the selected genes, while for polygenic gene expression evolution, the variance remains similar to neutral genes. Our computer simulations showed that this expectation from theoretical results (Jain & Stephan, 2015; Barton *et al*., 2017; Franssen *et al*., 2017) also holds for moderately sized populations and linked loci (Figure 2).

**Figure 6.**
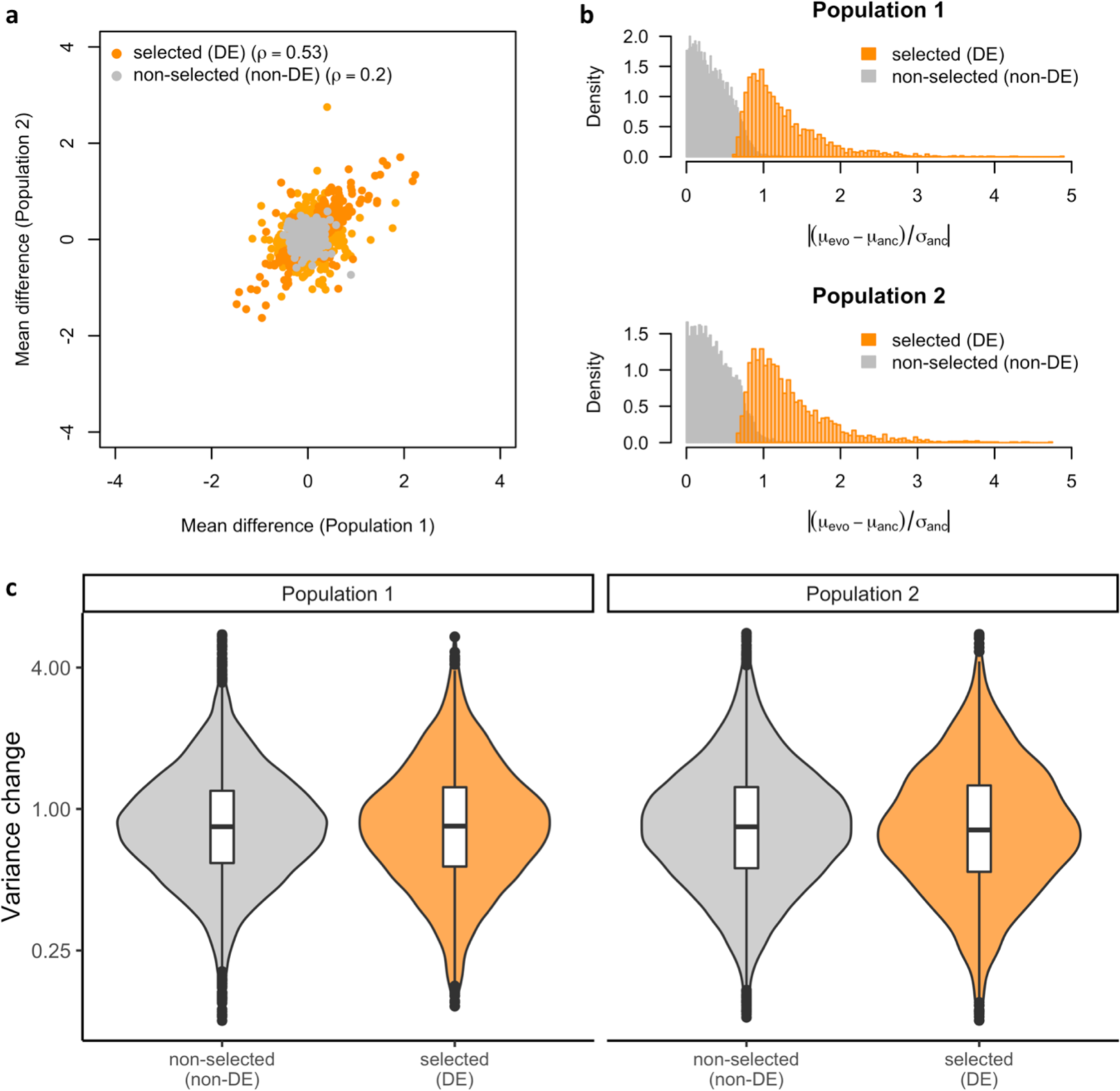
Evolution of phenotypic mean and variance over 100 generations of adaptation in empirical data. **a.** The evolution of gene expression means during adaptation in the two replicates. For the genes with significant changes (DE, in orange), the changes are correlated between replicates (Spearman’s rho = 0.53). For the genes without significant changes (non-DE, in grey), the correlation between replicates is much lower (Spearman’s rho = 0.2). **b.** The evolution of gene expression means scaled by the ancestral variation. For the DE genes (in orange), the median change is around one standard deviation of the ancestral expression value, suggesting mild shift in trait optimum in the novel environment. For the non-DE genes (in grey), the changes in expression are mostly negligible. The same pattern is seen in the second replicate. **c.** The change in expression variance during adaptation for DE and non-DE genes. In both replicates, the distribution of variance changes is indistinguishable between DE genes (orange) and non-DE genes (grey) (t-test, p > 0.05 for both replicates). The almost identical pattern is observed when the biological coefficient of variation (BCV; see materials and methods) is used to estimate the expression variance (Supplementary Figure S4).

The reduction in population size from a natural population to laboratory cultures would result in a continuous reduction in phenotypic variance over time even under neutrality, as shown in the simulations (Figure 2). Hence, a robust inference of the adaptive architecture of gene expression relies on the comparison of putative selected (DE) and neutral (non-DE) genes. In both replicates the expression variance of putative neutral (non-DE) genes dropped to 84% of the ancestral variance, which is quite close to the neutral cases in our simulations. While it would be desirable to test each putatively selected (DE) gene independently, the moderate sample size (∼20 per population) does not provide sufficient power to do so (Supplementary Figure S8). Rather, we considered all putatively selected (DE) genes jointly and compared their variance changes to the ones of putatively neutral (non-DE) genes. Remarkably, we find that the changes in variance of putatively selected genes with significant mean expression changes are indistinguishable from the genes that do not change their mean expression (t test, p-value > 0.05; Figure 6c). The magnitude of gene expression variance changes in both sets of genes (median F-value = 0.85 and 0.84 for DE and non-DE genes, respectively, in population 1, and 0.82 and 0.84 in population 2) is quite close to the neutral conditions in our computer simulations. This suggests that selection on mean expression did not significantly change the expression variance during adaptation. This is further supported by the lack of correlation between the evolutionary changes in gene expression variance and mean (Lai & Schlötterer, 2022). In our simulations we assumed that the traits are independent from each other, which may not be the case for gene expression data. Co-regulation (non-independence) among gene expression may affect our inference. We address this issue by using only 1000 genes with reduced correlation to diminish the impact of gene co-regulation for the analysis (see supplementary information and supplementary figure S10). Despite this down-sampling, we still observed no difference in variance change between DE and non-DE genes using this approach (Supplementary figure S11). Based on our analyses we suggest that variation in expression of most putatively adaptive genes has polygenic basis (i.e. more than five contributing loci).

Since we only explored two time points (F0 and F103) rather than a full time series, it may be possible that an oligogenic basis could also result in a similar phenotypic variance as neutral phenotypes like what is observed for a polygenic architecture (Supplementary Figure S9). This can be seen in an intuitive case when a single/few major effect allele(s) starts at a low frequency and becomes fixed (Yoo *et al*., 1980). Since an oligogenic basis results in a highly parallel genomic selection response (Supplementary Figure S12), it is possible to distinguish polygenic and oligogenic architectures with phenotypes from two time points only when genomic data are available. Because the genomic signature in the same experiment uncovered a highly heterogeneous selection response (Barghi *et al*., 2019), we can exclude the unlikely explanation of an oligogenic architecture resulting in a similar expression variance as non-selected genes. Rather, we suggest that the adaptive response in gene expression could be explained by a polygenic architecture (i.e. more than five contributing loci).

## Discussion

Population genetics has a long tradition of characterizing adaptation based on the genomic signature of selected loci (Nielsen, 2005). Nevertheless, for selected phenotypes with a polygenic architecture, the contribution of individual loci to phenotypic change may be too subtle to be detected with classic population genetic methods (Pritchard *et al*., 2010). Although identification of many small-effect loci is challenging, it may be possible to determine the number of contributing loci based on the dynamics of phenotypic variance.

Inspired by the Castle-Wright estimator (Castle, 1921) that estimates the number of loci contributing to a quantitative trait from the phenotypic variance of the F2, we propose that the temporal heterogeneity of the phenotypic variance can potentially be used to infer the number of loci contributing to the adaptive response of a phenotype as well as other parameters of the adaptive architecture. Reasoning that experimental evolution is probably the best approach to obtain phenotypic time series, we performed computer simulations specifically tailored to typical experimental evolution studies with *Drosophila*. We demonstrated that, in an experimental evolution setup, the temporal dynamics of the phenotypic variance is strongly affected by the number of contributing loci and other parameters of the adaptive architecture such as the distribution of effect size.

We acknowledge that the proposed approach for the characterization adaptive architecture critically depends on the power to distinguish the variance changes of selected traits from the neutral expectations. Since the accurate inference of gene expression variance is key, we partitioned the expression variance of each gene among three independent F1 families consist of almost identical, heterozygous individuals (see Supplementary Information). We show that on average, ∼50% of expression variance can be attributed to random error - a combination of technical (library preparation, RNA-extraction, sequencing) and biological (stochastic gene expression differences among individuals) noise. We included this error in our simulations by adopting the empirical heritability distribution (heritability of every gene inferred from this experiment) such that random errors relative to the ancestral phenotypic variance were introduced to the calculation of phenotypic values at each generation for each simulated trait. The heritability estimates in our experiment may be too conservative (i.e. too high) because we assumed no environmental heterogeneity. We also used heritability estimates from another gene expression study on *Drosophila* which included more lines and accounted for environmental heterogeneity (Ayroles et al., 2009). The results are qualitatively similar (Figure 3).

Another potential limitation of our analysis is that we assumed neutrality of non-DE genes, it may, however, be possible that they are also subject to stabilizing selection, but without shift in trait optimum. In this case, the same change in variance is expected for both groups of traits - even under an oligogenic architecture, as shown by computer simulations (Supplementary Figure S13a and b, see supplementary information). If stabilizing selection is also operating on non-DE genes, the change in variance should be correlated between replicate populations because each trait experiences the same strength of stabilizing selection (each trait has the same heritability in both replicates, but it differs between traits). This prediction was confirmed in our computer simulations (Supplementary Figure S13c). In our empirical data, however, the variance change was more strongly correlated between two evolution replicates for DE genes (r=0.07) than for non-DE genes (r=0.02) differ from each other, implying less selection for variance on non-DE genes. This difference in correlation is highly significant when we compared the correlation coefficient of DE genes to the same number of randomly drawn non-DE genes (p<0.01, Supplementary Figure S13d). Given that the distribution of correlation coefficients for non-DE genes is very close to the expectations under neutrality from computer simulations, we think that the behavior is better approximated by neutrality for most of the non-DE genes, rather than assuming that similar levels of stabilizing selection for both classes of genes. It is not clear, however, whether our results reflect a much broader fitness function determining the evolution of non-DE genes per se, or the simple laboratory environment relaxes selection on non-DE genes.

Given the limited power to infer the number of contributing loci for each expression phenotype (Supplementary figure S8), we grouped all putatively selected expression traits and limited our inference of the adaptive architecture to these genes jointly. This approach makes the implicit assumption that all expression traits in this group are independent of each other and have similar level of complexity in their adaptive architectures. Thus, the joint inference could potentially compromise the detection of a minor subset of genes whose expression evolution is under simple genetic control.

Our results may also provide some insights into the molecular basis of gene expression evolution. Gene expression is modified by many trans-acting factors and some cis-regulatory variation (Hill *et al*., 2021). Adaptive gene expression changes can be either driven by polymorphism in *cis*-regulatory elements or by *trans*-acting variants. While interspecific differences in gene expression are predominantly caused by *cis*-regulation (Wittkopp *et al*., 2004), intraspecific variation is mostly driven by trans regulatory changes (Wittkopp *et al*., 2008; Suvorov *et al*., 2013). Adaptive gene expression changes which are well-characterized on the molecular level typically have a *cis*-regulatory basis that is not only frequently associated with the insertion of a transposable element (e.g.: (Daborn *et al*., 2002)), but also sometimes with multiple regulatory variants (Endler *et al*., 2018). Two lines of evidence suggest that *cis*-regulatory variation cannot be the driver of adaptive gene expression changes observed in this study. First, the mutational target size is too small to harbor a sufficiently large number of alleles segregating in the founder population. Second, too few recombination events occur during the experiment to uncouple regulatory variants located on a given haplotype such that they could generate a signal of polygenic adaptation. More likely, the polygenic adaptive architecture of gene expression change reflects the joint effects of many *trans*-acting variants.

Because we could only analyze phenotypic data from two time points, the founder population and replicate populations evolved for 103 generations, we were not able to obtain a more quantitative estimate of the number of contributing loci, in particular as other parameters of the adaptive architecture are not known and need to be co-estimated. In addition, with only two time points, for a few parameter combinations, an oligogenic response can also result in a similar phenotypic variance change as a polygenic one but with a much higher parallel response of genomic markers (Supplementary Figure S12). Hence, not only more time points describing the phenotypic trajectory, but also some genomic data could contribute to inferring the adaptive architecture in experimental evolution studies.

The extension of the concept proposed in this study to natural populations could face several challenges that warrant extra caution and further investigation. First, the above-mentioned challenge of sample size would be more pronounced in a natural setup. Second, phenotypic time series over evolutionary relevant time scales are costly (but see (Clutton-Brock & Pemberton, 2004)) and third, the distinction of environmental heterogeneity from genetic changes is considerably more challenging than under controlled laboratory conditions.

## Materials and Methods

### Partitioning gene expression trait variance in three F1 families

Since the accurate measurement of gene expression variance is key to this study, we analyzed the expression variance for each gene among three independent F1 families obtained from crossing two highly inbred strains (see Supplementary Information and Table S1). We show that on average, ∼50% of expression variance can be attributed to random error - a combination of technical (library preparation, RNA-extraction, sequencing) and biological (stochastic gene expression differences among individuals) noise (Supplementary Figure S1). This confirms the importance to separate the biological and technical variance in the dataset, as we did in the empirical data analysis (see below).

### Computer simulations

We performed forward simulations with MimicrEE2 (Vlachos & Kofler, 2018) using the qff mode to illustrate the influence of the genetic architecture on the evolution of phenotypic variance during the adaptation to a new trait optimum (Figure 1a). With 189 founder haplotypes (Barghi *et al*., 2019), we simulated quantitative gene expression traits under the control of different numbers of loci (M = 5, 25, 50, 100, 200 and 1000) in a population with an effective population size of 300 (reflecting the estimated effective population size in an experimentally evolving population (Barghi *et al*., 2019)). We used the empirical linkage map from *Drosophila simulans* (Howie *et al*., 2019) to account for linkage. For each trait, we assume an additive model and a negative correlation (r = -0.7, reflecting the observed relationship from Otte *et al*. (2020)) between the effect size and starting frequency (Figure 1b). We used the *correlate()* function implemented in “fabricatr” R package (Blair *et al*., 2019) to generate the effect sizes. The distribution of the effect sizes of loci controlling a trait can vary depending on the shape parameter of gamma sampling process (shape = 0.5, 2.5 and 100, Figure 1c). The mean of the effect sizes from 3 different shape parameters was standardized to 1/M. As the focus of our study is on expression traits, we used the distribution of heritabilities obtained from empirical gene expression data (Ayroles et al. 2009 and this study, see supplementary information) to assign the environmental/technical variance to a simulated trait (see supplementary information; Supplementary Figure S1) (i.e. for each simulated trait, its heritability and associated environmental/technical variance are sampled from the empirical distribution). We do not simulate de novo mutations as they are not contributing to adaptation on this short time scale (Burke *et al*., 2010). To simulate stabilizing selection with trait optimum shift, we provided the Gaussian fitness functions with mean of 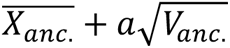 and standard deviation of 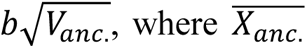 is the ancestral phenotypic mean and *V_anc_*. is the ancestral genetic variance (Figure 1a). Parameter “a” determines the distance of optimum shift, which is set to one (similar to the empirical case, Figure 6b) or three (following (Hayward & Sella, 2019)). Parameter “b” indicates the phenotypic constraint would be at trait optimum. The value 3.6 for parameter “b” was taken from Hayward and Sella (2019). In this study, we increase and decrease it by 50% to explore its impact (1.8 or 5.4). For the neutral case, we assumed no variation in fitness among all individuals. For each scenario, 1000 traits were simulated, and each trait was affected by a different set of loci. We compared the phenotypic variance of selected and neutral traits for each genetic architecture. For each trait under each scenario, the phenotypic variance was estimated at different generations and compared to the ancestral phenotypic variance at generation 1 to illustrate the dynamic of phenotypic variance during the evolution 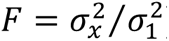, where x stands for the number of generations. We note that we do not assume that the ancestral population has reached an equilibrium because the ancestral population in a typical experimental evolution study is often phenotyped in the new environment.

In addition, we evaluated how different experimental designs (e.g.: different population sizes in the experiment) affect the ability to discriminate between different adaptive architectures. We simulated the evolution of traits controlled by a different number of loci (M = 5, 25, 50, 100, 200 and 1000) with varying effect sizes (shape parameter of gamma sampling process = 2.5) using a Gaussian fitness function with a mean of 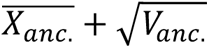 and a standard deviation of 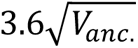 with population size = 1200.

### Evaluating the ability to distinguish adaptive architectures based on trait variance evolution

We propose that the temporal dynamics of the phenotypic variance of selected traits can be exploited to characterize the genetic basis of adaptation. This concept could be applied either to a single trait under selection or to a group of selected traits. Depending on whether a single trait or a group of traits is studied, different research objectives can be pursued. Generally speaking, a more complex architecture is expected to result in a variance pattern that resembles neutral evolution (i.e. no change in variance). A lower bound for the number of contributing loci can be inferred by asking for the maximum number of contributing loci causing a variance change during evolution, which significantly deviates from neutral expectations. With increasing uncertainty about the key parameters, the precision of this approach is reduced. Given that many parameters are typically not known, we performed computer simulations for different numbers of loci with a distribution of effect sizes and asked for each focal parameter combination whether a significant deviation from neutral expectations was observed. The number of simulation runs, which deviate from neutral expectations for *x* loci is used as power estimate to reject the null hypothesis of no difference to the variance under neutrality under a given architecture and sample size. In other words, it reflects the confidence to exclude an adaptive architecture with *x* or fewer contributing loci when we observed no significant difference in variance.

#### Power estimates for the analysis of a single trait

For each simulated trait under different genetic control, we tested whether the variance of this trait changed more than expected under neutrality (F of 0.9 according to the neutral simulations) after 100 generations of selection. First, we assumed that all simulated 300 individuals were phenotyped at both time points. For the 1000 traits with a given genetic architecture (*x* contributing loci) under each selection scenario, we calculated the power to reject the null hypothesis of an adaptive architecture no difference to the variance under neutrality under a given architecture and sample size. Since it is not possible to phenotype all individuals, we also investigated whether and how a reduced sample size compromises the power to reject an adaptive architecture of a single trait based on the change in variance. We randomly selected 20 individuals from each simulation run to estimate the variance and tested whether the magnitude of variance change for a given trait under selection is significantly different from the neutral expectation.

#### Power estimates for the analysis on a group of traits

We assessed the power to infer the adaptive architecture for a group of selected traits under the assumption that most traits of interest have similar level of complexity. In this case, a test on the average variance changes (F) between a group of selected traits and another group of neutral traits is required. We simulated 1,000 independent selected and 1,000 neutral traits for each genetic architecture and selection regime, and estimated the variance before and after 100 generations of evolution for the selected and neutral traits. For each trait, we measured the phenotypic variance for a sample of 20 individuals and we performed a t-test to compare the distribution of variance change (F) between selected and neutral traits. The power was determined by 100 iterations of sampling 20 individuals from a simulated population of 300 individuals.

We further evaluated how different experimental designs affect the power to detect simpler adaptive architectures. We used the data from the scenario with shape parameter 2.5 and Gaussian fitness functions with mean of 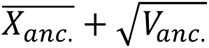 and standard deviation of 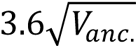. With this dataset, we calculated the power for different sample sizes (n=20, 30, 50, 70), generation times (gen=25, 50, 100, 200) and population sizes (N=300 and 1200).

### Experimental evolution

The setup of populations and evolution experiment have been described by Barghi *et al*. (2019). Briefly, ten outbred *Drosophila simulans* populations seeded from 202 isofemale lines were exposed to a laboratory experiment at 28/18 °C with 12hr light/12hr dark photoperiod for more than 100 generations. Each replicate consisted of 1000 to 1250 adults for each generation.

### Common garden experiment

The collection of samples from the evolution experiment for RNA-Seq was preceded by two generations of common garden (CGE). The common garden experiment was performed at generation 103 of the evolution in the hot environment and this CGE has been described in (Barghi *et al*., 2019; Hsu *et al*., 2019, 2020; Jakšić *et al*., 2020; Lai & Schlötterer, 2022). In brief, an ancestral population was reconstituted by pooling five mated females from 184 founder isofemale lines (Nouhaud *et al*., 2016). No significant allele frequency differences are expected between the reconstituted ancestral populations and the original ancestral populations initiating the experiment (Nouhaud *et al*., 2016). Furthermore, we do not anticipate that deleterious alleles acquired during the maintenance of the isofemale lines had a major impact on the phenotypic variance in the reconstituted ancestral population. The reason is that novel deleterious mutations occurring during the maintenance of the isofemale lines are present in a single isofemale line only. Given the large number of isofemale lines (184), such deleterious alleles occur in a low frequency in the reconstituted population with a small influence on the phenotypic variance (Walsh & Lynch, 2018). Furthermore, most of these deleterious alleles are present in heterozygous individuals and masked because deleterious alleles tend to be recessive (Charlesworth & Charlesworth, 2010). As described previously (Lai & Schlötterer, 2022), two replicates of the reconstituted ancestral population and two independently evolved populations at generation 103 were reared for two generations with controlled egg density (400 eggs/bottle) at the same temperature regime as in the evolution experiment. Freshly eclosed flies were transferred onto new food for mating. Sexes were separated under CO_2_ anesthesia at day 3 after eclosure, left to recover from CO_2_ for two days, and at the age of five days whole-body mated flies of each sex were snap-frozen at 2pm in liquid nitrogen and stored at -80°C until RNA extraction. More than 30 individual male flies from two reconstituted ancestral populations (replicate no. 27 and no. 28) and two evolved populations (replicate no. 4 and no. 9) were subjected to RNA-Seq. The protocols of RNA extraction and library preparation are described in (Lai and Schlötterer 2022).

### RNA-Seq data analysis for mean and variance evolution

The processed RNA-Seq data were obtained from Lai and Schlötterer (2022). To characterize the adaptive architecture of gene expression evolution, we compared the evolutionary dynamics of the variance between the genes with significant mean change and those without. The underlying assumption is that genes with significant mean expression changes are under selection and the rest of the transcriptome is not responding to the new environment (neutral). The genes with/without significant mean evolution for two evolved populations were taken from Lai and Schlötterer (2022).

We quantified the variance changes during adaptation for each gene by using the variance estimates of the expression (logCPM) of each gene in each population from Lai and Schlötterer (2022). Briefly, raw read counts of each gene were normalized with the TMM method implemented in edgeR. We then applied natural log transformation to the expression of each gene (counts per million (CPM)) to fit normal assumption for all subsequent analyses and make mean and variance independent from each other.

Due to the moderate sample size, we performed additional checks for the uncertainty of variance estimates. Jackknifing was applied to measure the uncertainty of estimator (Fukunaga & Hummels, 1989). The procedure was conducted independently on four populations, and we calculated the 95% confidence interval of the estimated variance (Supplementary Figure S2). Additionally, the robustness of variance estimation was also supported by the high correlation (rho = 0.79, Spearman’s rank correlation) in variance estimates between the two independently reconstituted ancestral populations. The change of gene expression variance was determined by the F statistics calculated as the ratio between the variance within the ancestral population and the variance within the evolved population of each gene. To test whether selection on mean expression generally alters the expression variance, we compared the F statistics of genes with significant changes in mean expression to the genes without.

We note that the natural log transformation (as in Lai and Schlötterer 2022) does not attempt to quantify and partition the noise due to the read sampling process at the gene level from the across-individual variance. Since we were primarily interested in the latter, we evaluated the effect of read sampling noise. We did an additional check by distinguishing the true biological variation and the measurement error ([eq1], from User’s Guide of edgeR) of each gene using the statistical method implemented in edgeR, where the true variance across individuals (biological coefficient of variation (BCV)) of each gene can be estimated (tag-wised dispersion).

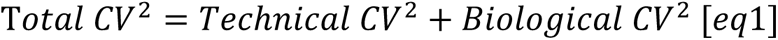

The proportion of technical CV^2^ in total CV^2^ was calculated. We showed that the average proportion of technical CV^2^ gradually decreased with increasing sample size (Supplementary Figure S3), and for our sample size (n = 22) it is 17%. The correlation between BCV^2^ and the variance estimates after log transformation is 0.98 (Spearman’s rho), suggesting that the two estimates are similar. As a sanity check, we repeated the comparisons of the variance changes in selected (DE) and neutral (non-DE) genes using BCV^2^. Similarly, the changes in variance of putative adaptive genes are indistinguishable from the genes that do not change their mean expression (t-test, p-value > 0.05; Supplementary Figure S3).

## Acknowledgments

Special thanks to David Houle, who provided fantastic support during the collection and establishment of the isofemale lines in Florida. We thank all member of the Institut für Populationsgenetik for discussion. We are grateful to Reinhard Bürger, David Houle, Dagný Ásta Rúnarsdóttir and anonymous reviewers for helpful comments on earlier versions of the manuscript. Neda Barghi, François Mallard and Kathrin Otte contributed to the common garden experiment. Illumina sequencing was performed at the VBCF NGS Unit (www.vbcf.ac.at). This work was support by the Austrian Science Funds (FWF, W1225) and the European Research Council (ERC, ArchAdapt).

## Author contribution

W.Y.L and C.S. conceived the study. V.N. prepared all RNA-Seq and supervised the maintenance of the evolution experiment. A.M.J supervised the common garden experiment. W.Y.L performed the simulation and data analysis. W.Y.L. and C.S. wrote the manuscript.

## Data accessibility statement

All sequencing data will be available in European Nucleotide Archive (ENA) under the accession number PRJEB37011 upon publication.

## Competing interests

The authors declare no competing interests.

**Correspondence and requests for materials** should be addressed to C.S.

