## Supplementary information for "Evolution of phenotypic variance provides insights into the genetic basis of adaptation"

**Supplementary text**

Estimating genetic variance in gene expression across F1 families

In this study, we included an experiment to evaluate how much of the expression variance can be explained by genetic variation.  Using RNA-Seq on individual flies, with 3-4 individuals each from three iso-female lines maintained at the same density and culturing conditions we performed PCA and calculated broad sense heritabilities (Supplementary file 1).

Six out of 184 founder iso-female lines from the evolution experiment and were maintained for one generation with controlled egg density (400 eggs/bottle) in the same environment as the main experiment (12h 28°C with light followed by 12h 18 °C with dark conditions). Using the offspring, we generated three crosses between two of the six lines each: FL 138 x FL 137, FL 157 x FL 112, FL 123 x FL 127: we combined 50 virgin females from one of the lines with 50 males from the other line, let them lay eggs under density control as above and maintained and froze their F1 offspring in the same way as in the main CGE: sexes were separated after mating at the age of three days and snap-frozen at the age of five days at 2pm. From each cross, we used four F1 males to prepare individual RNA-Seq libraries as described above.

Assuming no environmental heterogeneity, we decomposed the total variance of the expression of each gene measured in these individuals into the genetic difference among three different F1 families and environmental/technical error. The data were analyzed as follows:

Natural log-transformation was applied to CPMs of all genes to improve data normality (Rocke and Durbin 2003). We tested for the genetic variance of each gene separately using analysis of variance (ANOVA):

$$y_{ij} = \mu + \tau_i + \varepsilon_{ij},$$

Where i=1, 2, 3 (the $i^{th}$ families); j =1, 2, 3, 4 (the $j^{th}$ individuals in each cross). $y_{ij}$ is the observed expression level of a gene in a given sample, $\mu$ is the overall mean; $\tau_i$ is the effect of genetic background and $\varepsilon_{ij}$ is the random noise. We calculated the proportion of total variance explained by random error using the following equation:

$$variance\ explained\ by\ random\ error = \frac{sum\ squares\ of\ error\ (SSE)}{sum\ squares\ of\ total\ (SST)}$$

We found that on average, around 50% of the gene expression variance can be explained by the genetic difference among the three F1 families (Supplementary Figure S1). This implies that a substantial within-population (between individuals) gene expression variance in our focal experiment is contributed by the genetic components. Because we only used offspring from single vials, we may have overestimated the heritability if the environment in the vials differs. Nevertheless, since our heritability estimates are very similar to previous ones (Ayroles *et al.* 2009), we consider our estimates reliable.

The technical error will go into residual in this analysis. Since variance of lowly expressed genes are mainly dominated by technical error, we defined a cut-off of expression value for a more reliable variance estimate. Genes were binned based on their average expression value (lnCPM) which ranged from -0.8 to 4.1, by bin size of 0.1. The average proportion of variance explained by random error of each bin was calculated. The expression variance of genes with less than 1 count per million bases (CPM) is dominated by technical errors (Supplementary Figure S9). Thus, genes with less than 1 count per million base (CPM) were excluded for subsequent analysis.

<u>Down sampling process for gene expression data set</u>

Co-regulation (non-independence) among gene expression may affect our inference due to pseudo-replication. We address this issue by using only less-correlated genes for the empirical comparison. For all 10,583 expressed genes, we first calculated their similarity on the expression level (pairwise Pearson's correlation among the expression of genes). Based on these measurements, we performed hierarchical clustering to group the genes that are potentially co-regulated. 1,000 groups are identified (Supplementary Figure S10A). From each group, we randomly pick a gene so that we have 1,000 genes that are significantly less-correlated compared to the original gene set (KS test, p-value<2.2e-16; Supplementary Figure S10B). These 1,000 genes were further used for the comparison of variance change between DE and non-DE genes.

<u>Simulating stabilizing selection without shift in trait optimum</u>

It is possible that non-DE genes are subject to stabilizing selection without shift in trait optimum rather than neutrality. Hence, we performed additional computer simulations to evaluate the case under stabilizing selection without shift in trait optimum. We followed the parameter setting from the scenario with shape parameter 2.5 but the Gaussian fitness functions was changed into mean of $\overline{X_{anc.}} + 0\sqrt{V_{anc.}}$ and standard deviation of $3.6\sqrt{V_{anc.}}$. We evaluate the power to distinguish the variance changes under stabilizing selection with and without shift in trait optimum with different numbers of contributing loci (M = 5, 25, 50, 100, 200 and 1000). We also investigate the correlation of variance changes between evolution replicates in the three different scenarios (stabilizing selection with shift in trait optimum, stabilizing selection without shift in trait optimum, and neutrality). When stabilizing selection operates, the change in variance should be correlated between replicate populations but such correlation is not expected under neutrality.

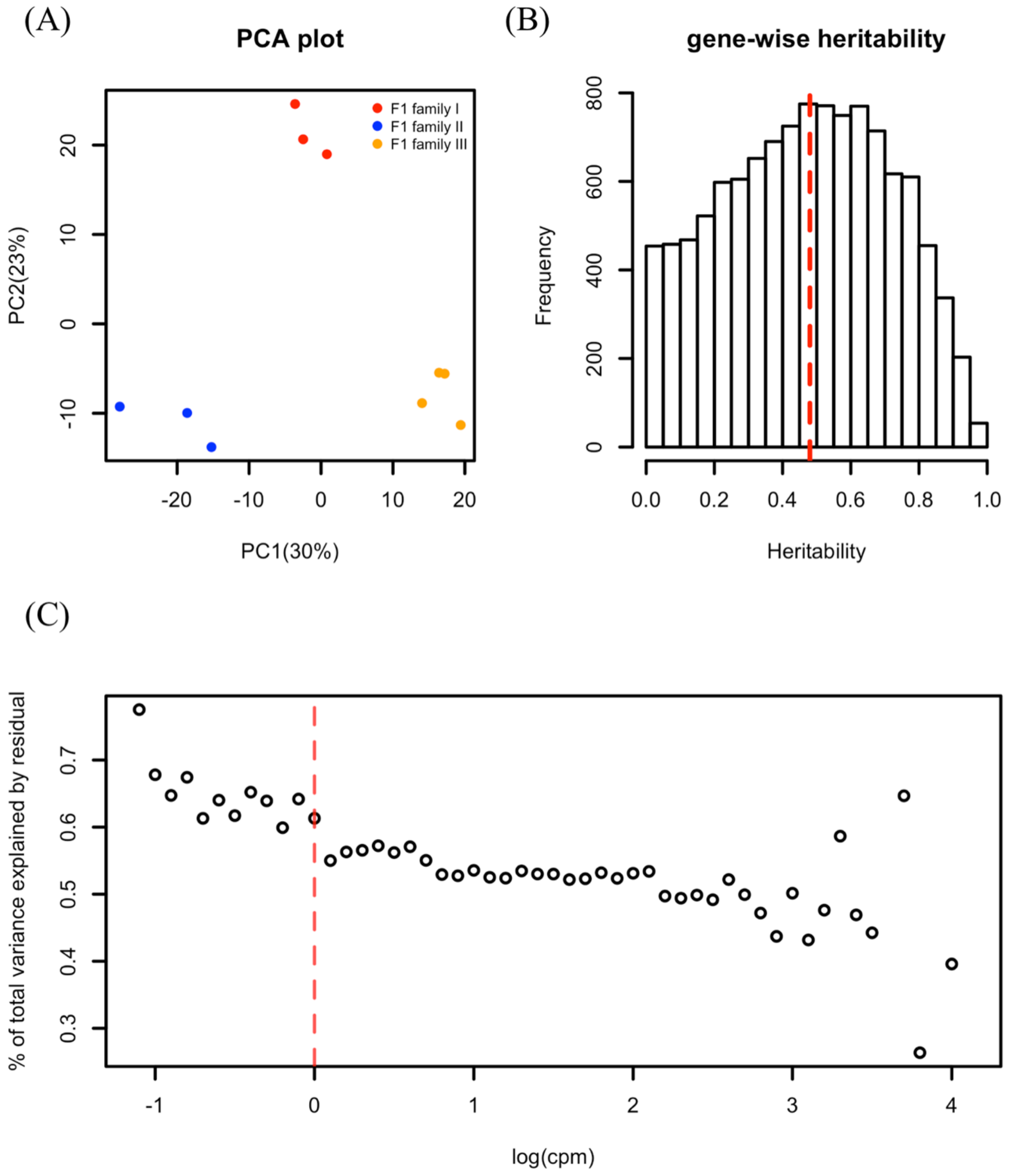


**Supplementary Figure S1. Genetic variance in gene expression across F1 families. (A)** Principal component analysis (PCA) on the transcriptomes of F1 individuals from three different crosses between the founder iso-female lines. Individuals from different families clustered nicely based on the first two PCs. (**B)** Gene-wise heritability on the transcriptomes of F1 individuals. On average, 48.8% of the total variance in gene expression was explained by the genetic difference between the individuals. (**C)** Genes were binned based on their average expression value (lnCPM) which ranged from -0.8 to 4.1, by bin size of 0.1. The average proportion of variance explained by random error of each bin was visualized. The expression variance of genes with less than 1 count per million bases (CPM) is dominated by residuals.

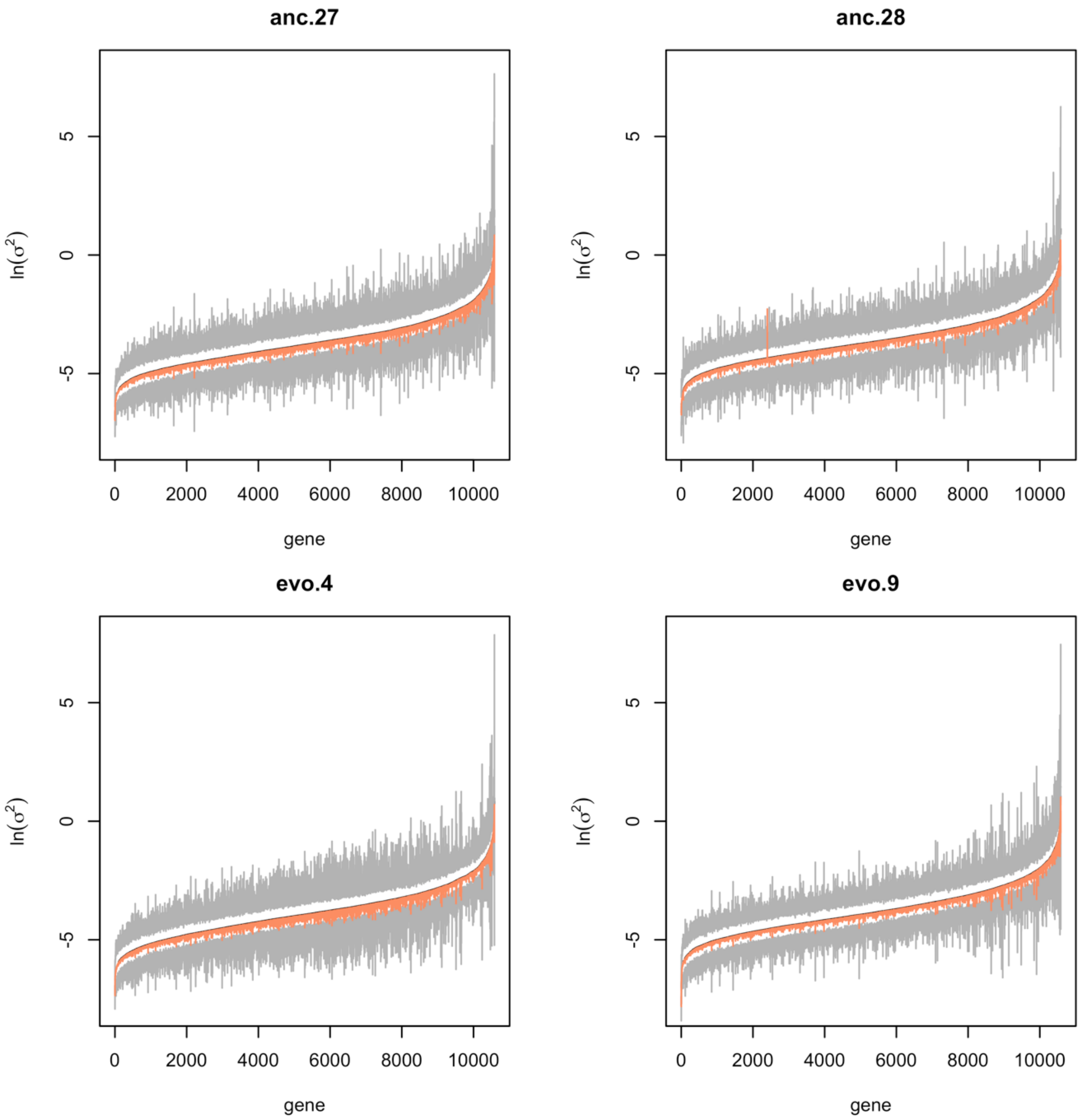


**Supplementary Figure S2. Robustness of the variance estimation using individual sequencing data.** Jackknife method was applied to measure the uncertainty of variance estimation within each population. Given a sample size of K, the procedure is to estimate the variance of each gene for K times, each time leaving one sample out. The procedure was conducted independently on 4 populations (anc.27, anc.28, evo.4 and evo.9). In each panel, we visualize Jackknife approximated 95% confidence interval for the variance estimates of each gene. The genes are ordered based on the average variance estimates (black dash line) on the x-axis. The upper and lower limits of the 95% confidence interval are indicated with grey curves. The salmon line denotes the observed value of the variance estimates. In most cases, the estimates lie in the confidence interval, suggesting robust estimation.

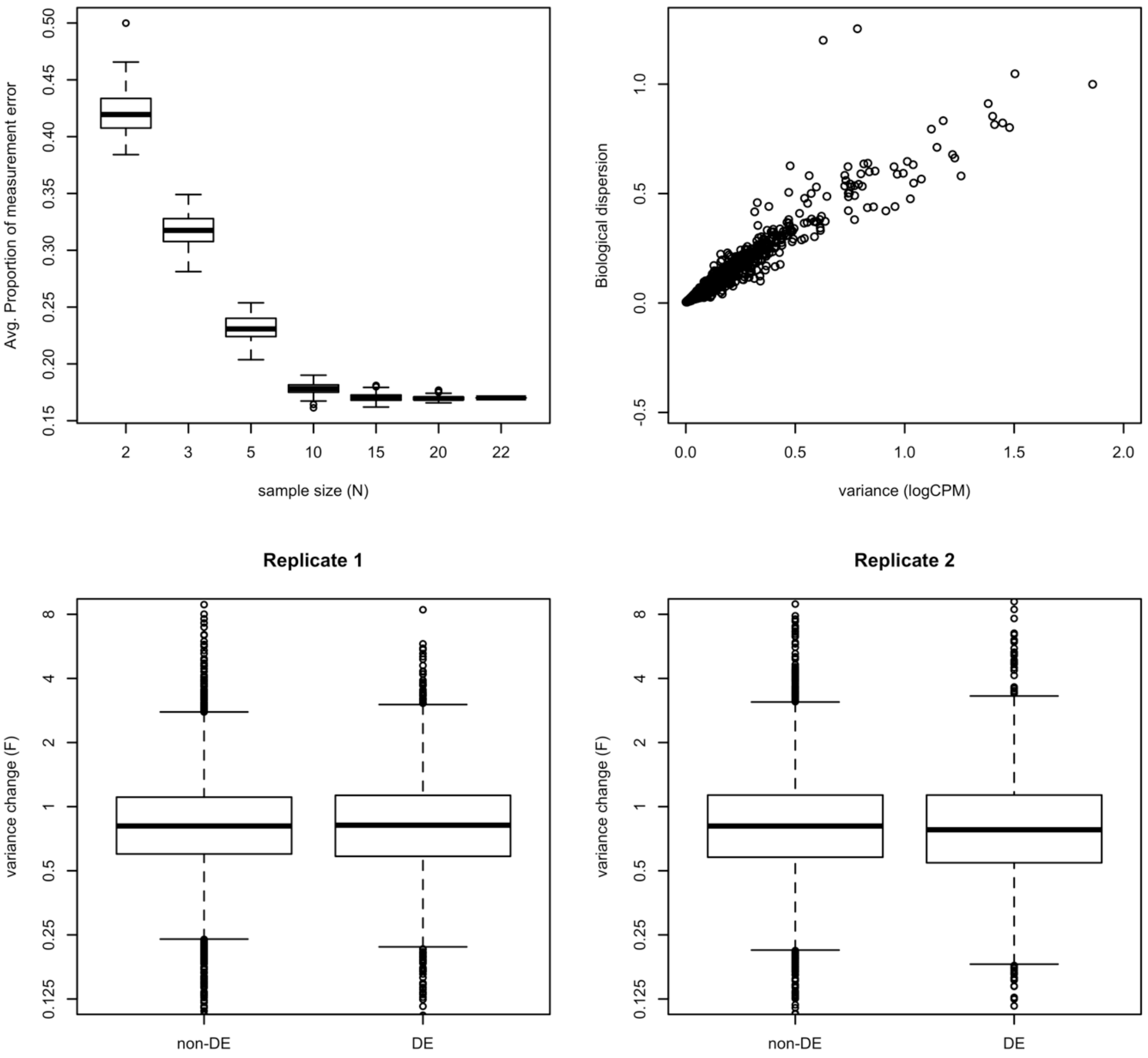


**Supplementary Figure S3. The decomposition of measurement error in RNA-Seq data and its implication for the tests of variance evolution. (A)** The relationship between the average proportion of measurement error among the RNA-Seq samples and the sample sizes is visualized. As the sample size increases, the measurement error of RNA-Seq data is well controlled. The results are consistent across replicate populations. For simplicity, only the result in the first replicate of ancestral population is shown. **(B)** We observed a strong correlation between the variance estimates after log-transformation and the biological coefficient of variation (BCV) estimated from edgeR, which claims to exclude the measurement error. Such strong correlation suggests that measurement error in RNA-Seq data wouldn’t have strong impact on the test of variance of evolution. The results are consistent across replicate populations. For simplicity, only the result in the first replicate of ancestral population is shown. **(C)** Using the BCV estimates, we repeated the comparison between the variance changes ($F = \frac{BCV_{evo}^2}{BCV_{anc}^2}$) of the genes with and without significant evolution in mean expression. In both replicates, the distribution of variance changes is indistinguishable between DE genes and non-DE genes (t-test, $p > 0.05$ for both replicates).

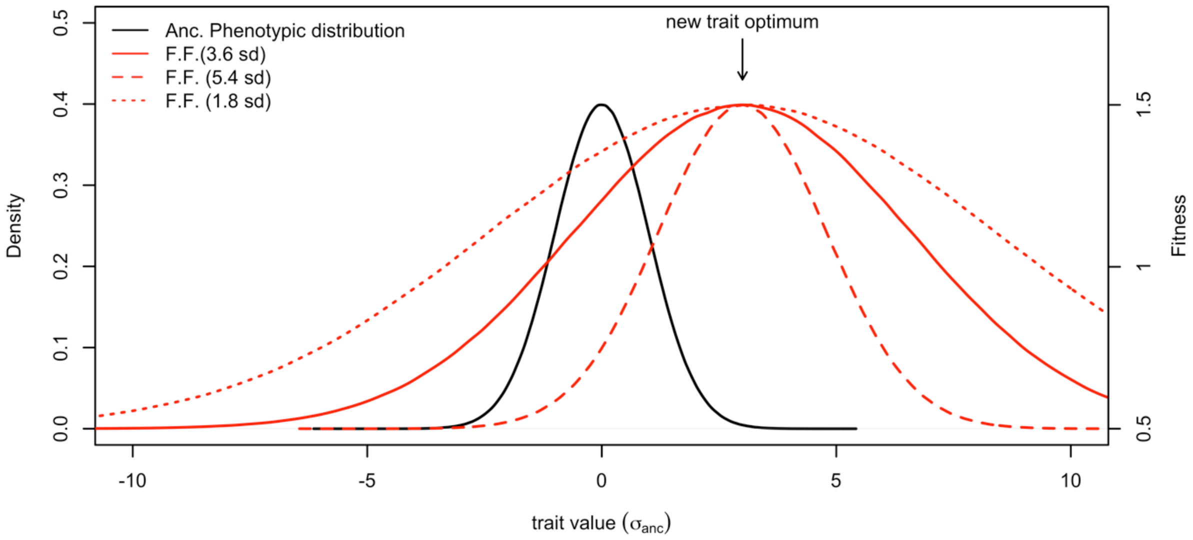


**Supplementary Figure S4. The evolutionary scenario for distant optimum shift.** We consider the case when a quantitative trait (in black) experiences a sudden shift in trait optimum under stabilizing selection. The imposed fitness functions (F.F.) are illustrated in red. The new trait optimum is set away from the ancestral trait mean by three standard deviation of the ancestral trait distribution for distal shift. To vary the strength of stabilizing selection, the variance of the fitness function is set as 1.8, 3.6 and 5.4 standard deviation of the ancestral trait distribution.

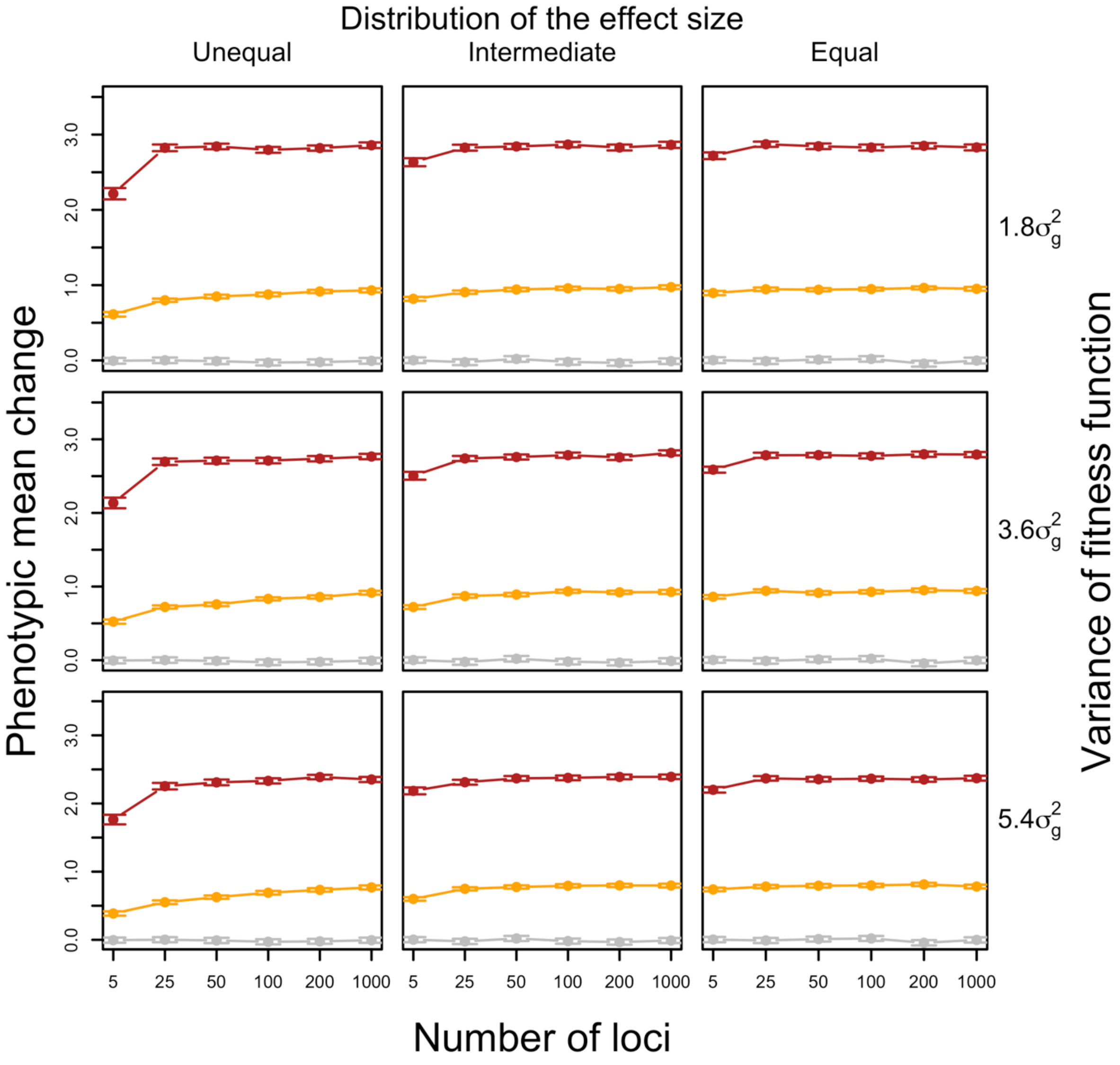


**Supplementary Figure S5. The changes in phenotypic mean when adapting to optimum shift.** The changes in phenotypic mean after 100 generations adapting to a mild/distant optimum shift (orange/red) are compared to the changes under neutrality (grey) on y axis. The changes in phenotypic mean are scaled by the standard deviation of the ancestral trait distribution. The simulations cover traits controlled by varying numbers of loci underlying the adaptation (x axes) with three different distributions of effect sizes (columns) and under different strength of stabilizing selection (rows). For each scenario, 1000 traits have been performed. In most cases, the traits under selection (orange/red) shift their means by one/three standard deviation of the ancestral trait distribution (i.e. reaching the new trait optimum) while the neutral traits (grey) stay unchanged. The error bar indicates the 95% confidence interval for 1000 simulated traits.

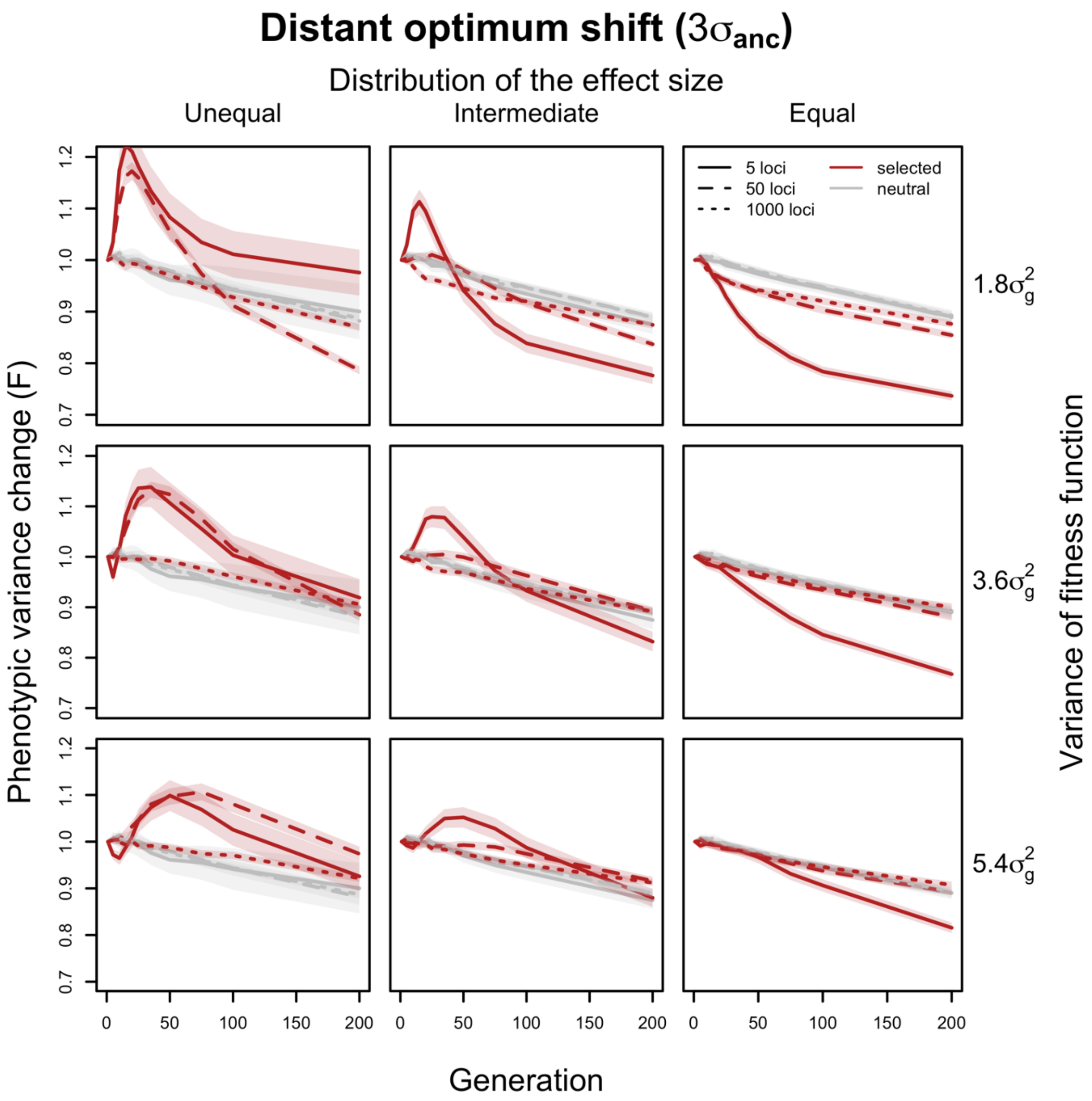


**Supplementary Figure S6. The trajectory of expected changes in phenotypic variance when adapting to a distant optimum shift.** The changes in phenotypic variance within 200 generations adapting to a distant optimum shift (red) are compared to the changes under neutrality (grey) on y axis. The average change in variance among 1000 traits (F) is calculated as the ratio of phenotypic variance between each evolved time point (generation x) and the ancestral state ($\sigma_x^2/\sigma_1^2$). The translucent band indicates the 95% confidence interval for 1000 simulated traits. The simulations cover traits controlled by varying numbers of loci underlying the adaptation with three different distributions of effect sizes (columns) and under different strength of stabilizing selection (rows). For each scenario, we simulated 1000 traits. Only traits with the most (dotted lines, 1000 loci), intermediate (dash lines, 50 loci) and the least (solid lines, 5 loci) polygenic architectures are shown. Unlike the continuous decreasing pattern in the cases with mild optimum shifts, the variance of traits controlled by a few loci (5 loci) with largely dispersed effects would increase first and then decrease when the effect sizes of contributing loci are dispersed (red solid lines). Nevertheless, for traits with extremely polygenic basis, the phenotypic variance always stays stable over time (red dotted lines).

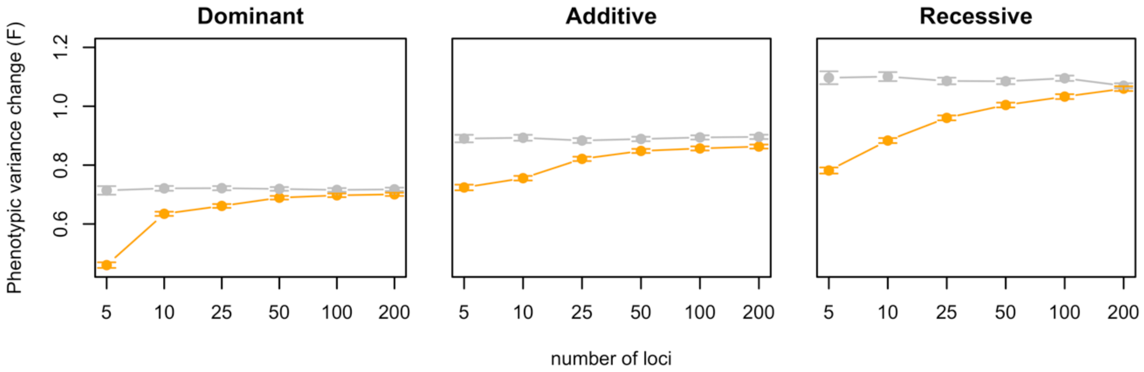


**Supplementary Figure S7. The expected changes in phenotypic variance for traits controlled by dominant and recessive alleles.** The changes in phenotypic variance after 100 generations adapting to a mild optimum shift (orange) are compared to the changes under neutrality (grey) on y axis. The change in variance (F) is calculated as the ratio between the evolved and ancestral phenotypic variance ($\sigma_{100}^2/\sigma_0^2$). This simulation covers traits controlled by varying numbers of loci underlying the adaptation (x axes) with recessive, additive and dominant effects. For each scenario, we simulated 1000 traits. No matter how the dominance varies, the variance of the trait decreases drastically when the adaptation is controlled by a small number of loci. As the number of contributing loci increases, the phenotypic variance becomes more stable. The error bar indicates the 95% confidence interval for 1000 simulated traits.

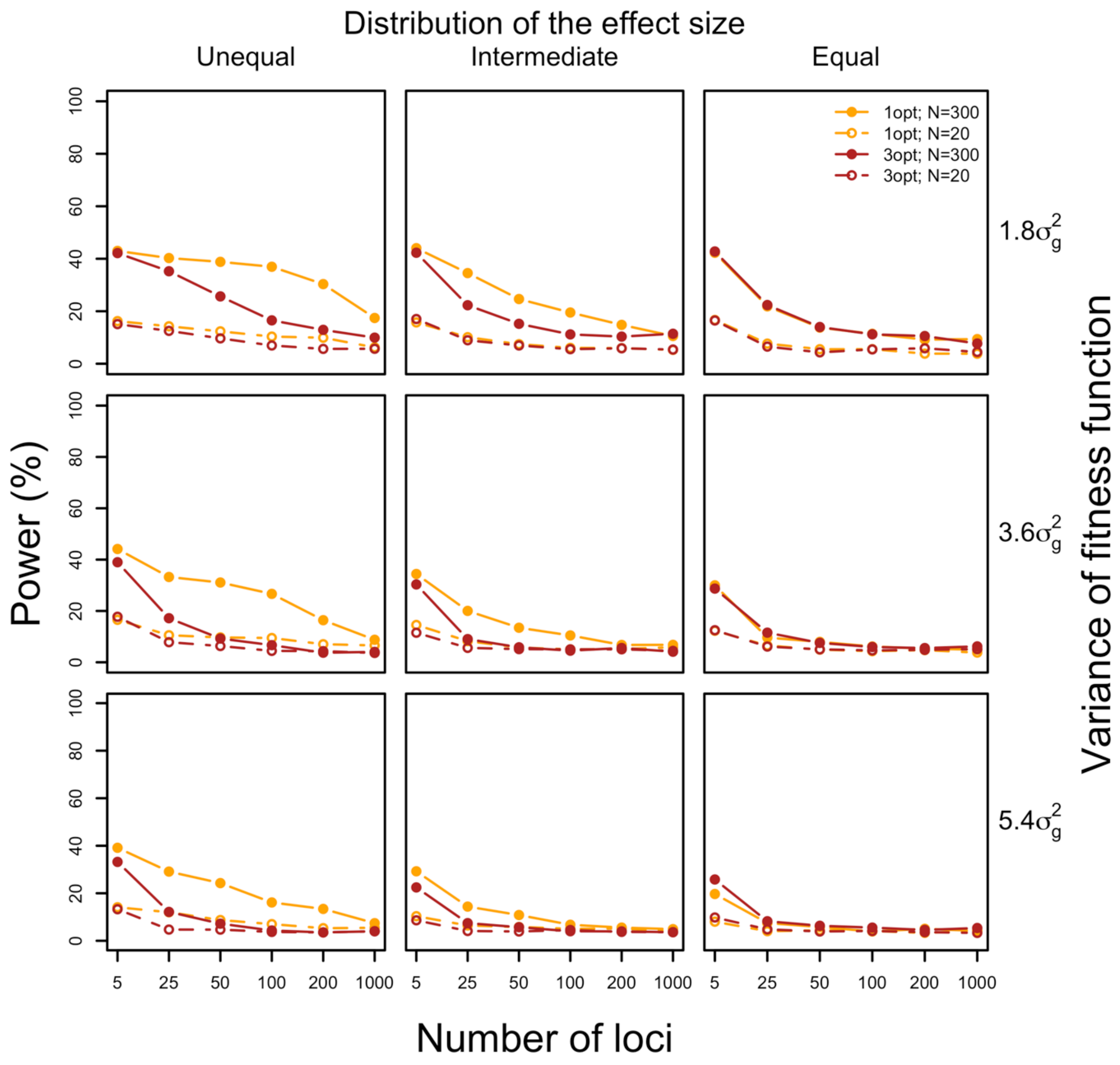


**Supplementary Figure S8. Power for detecting significant variance changes for a single trait.** With each genetic architecture (x-axis) under each selection scenario, we calculated how often we can detect a significant difference between the variance change of a single trait after 100 generations of selection and the neutral expectation (F of 0.9 according to the neutral simulations) (y-axis). The power is largely limited by the sample size. When all 300 individuals were considered, only 40% of power is observed under simple genetic control (5 loci). When the sampling processes of 20 individuals from the 300 simulated individuals was considered, there is nearly no power for the inference.

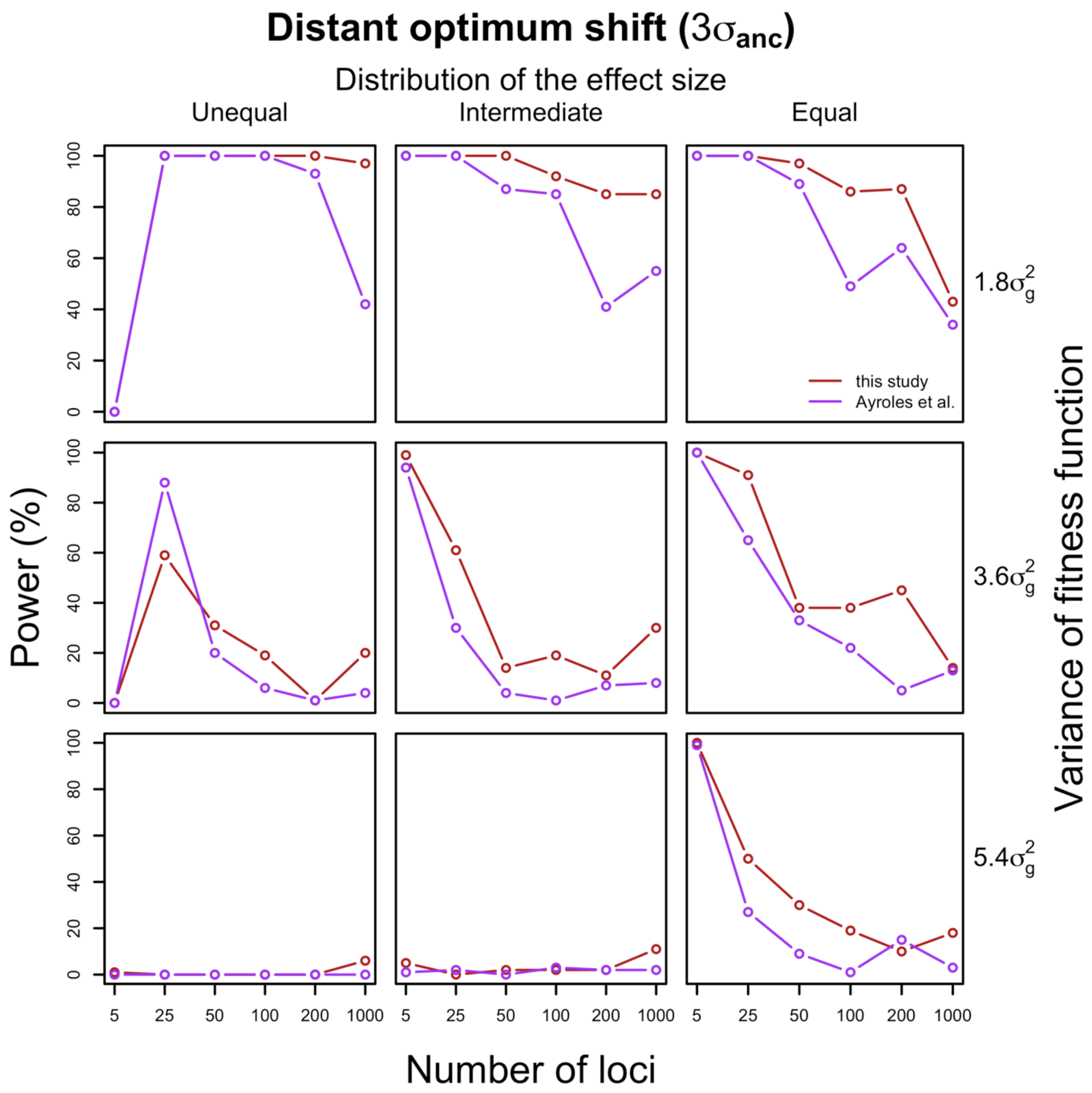


**Supplementary Figure S9. Power for detecting significant different variance change between a group of selected and neutral traits under distant optimum shift.** For each set of the 1000 traits controlled by different numbers of loci (x-axis) with varying effect sizes (columns) under each selection strength (rows), we calculated how often the variance changes of these 1000 selected traits differs from 1000 neutral traits after 100 generations (y-axis). The same genetic architecture is assumed for all 1000 selected traits with with a sample size of 20. In five out of the nine parameter combinations, we have nearly 100% power to detect a significant difference when all traits were under simple genetic control (5 loci). The power gradually decreases when the number of loci increased. Nevertheless, in the other parameter combinations, exception can be found for the traits controlled by a few loci (5 loci) with largely dispersed effects (see Discussion).

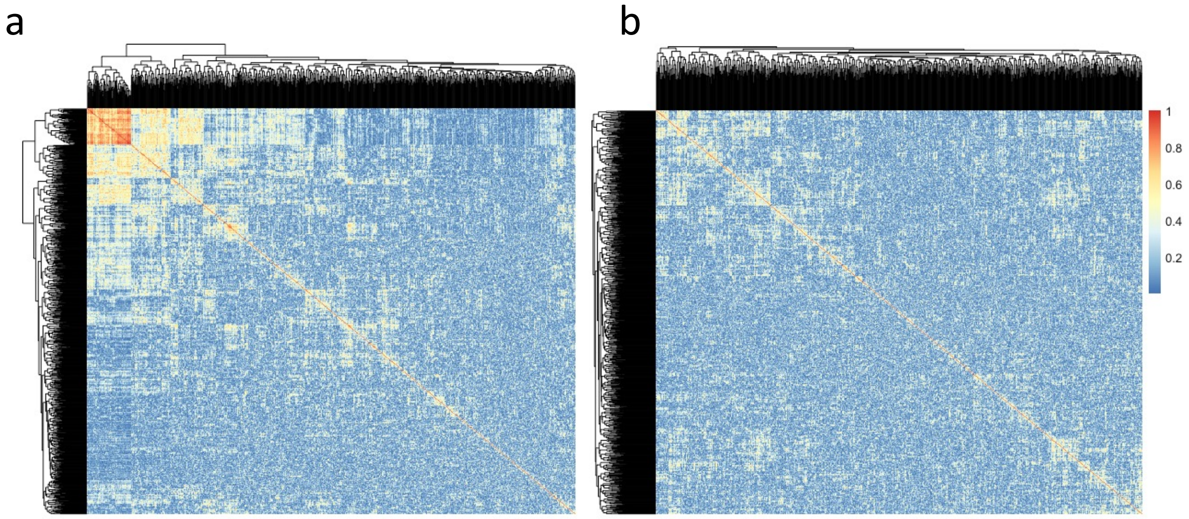


**Supplementary Figure S10. Pair-wise Pearson's correlation coefficient of the expression value between a. all genes b. down-sampled genes.** After down-sampling, the expression among genes became significantly less correlated with each other (KS test, p-value<2.2e-16).

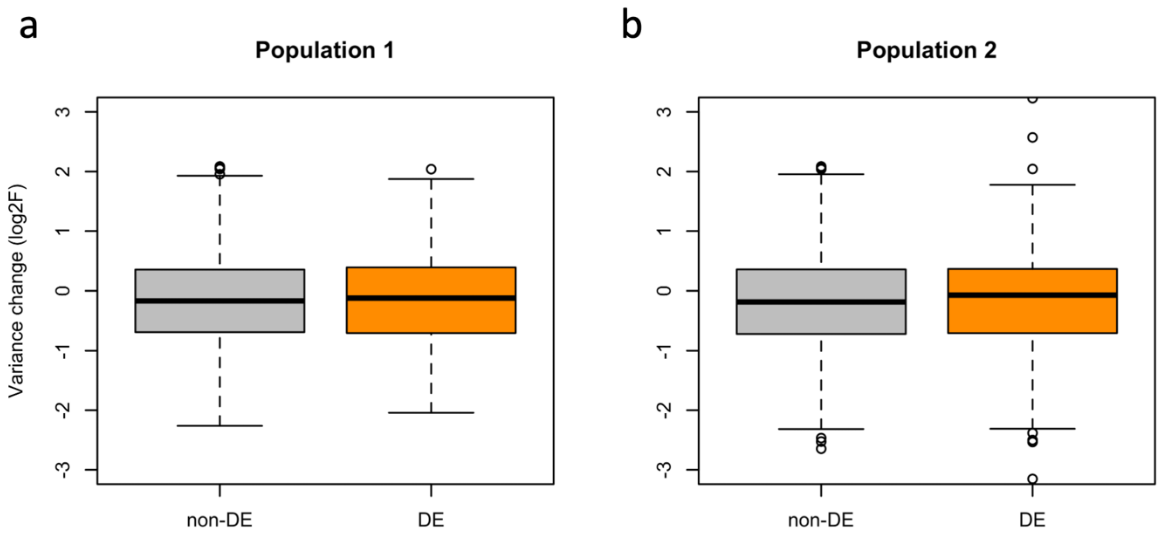


**Supplementary Figure S11. The change in expression variance during adaptation for DE and non-DE genes (down sampled 1000 gene set).** In both populations, the distribution of variance changes is indistinguishable between DE genes (orange) and non-DE genes (grey) (t-test, $p > 0.05$ for both replicates) for less correlated 1,000 gene set.

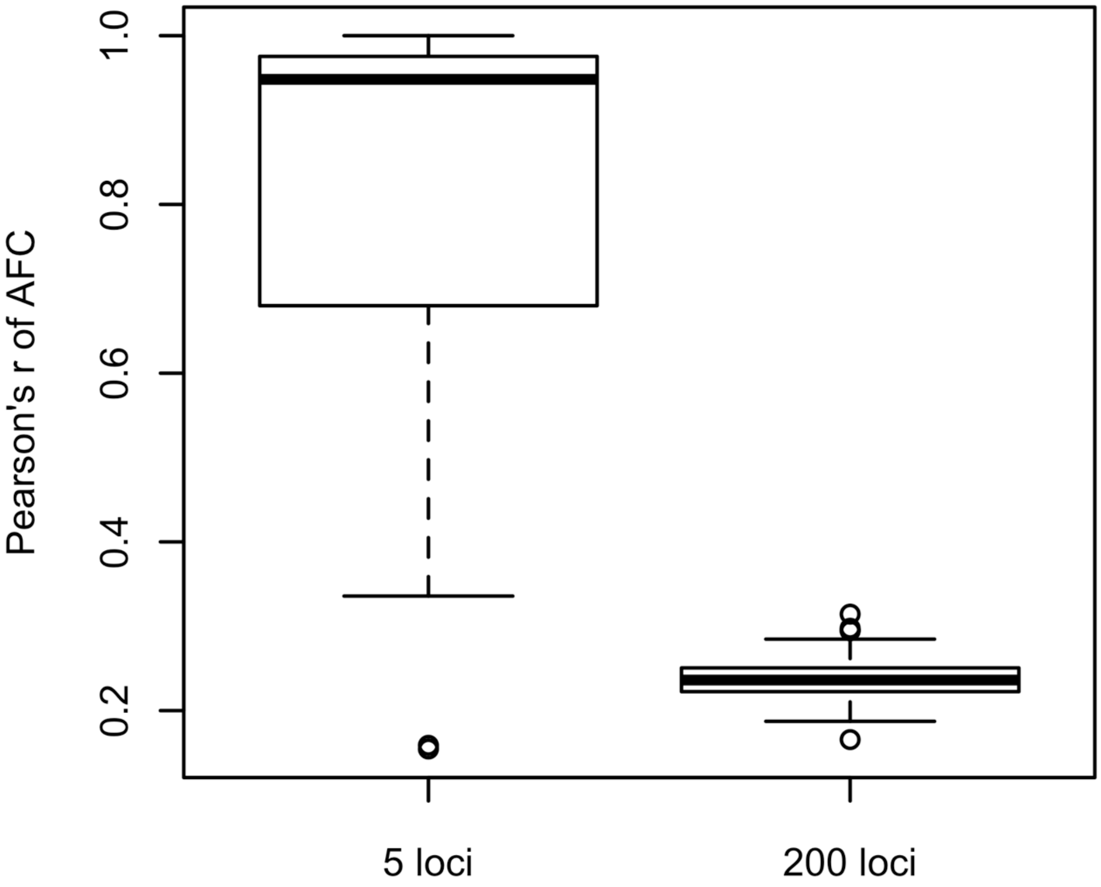


**Supplementary Figure S12. Parallelism in genomic response across 10 evolved replicates for traits under the control of five loci and 100 loci.** For the loci with allele frequency change of at least 10% in 100 generations, the average Pearson's correlation coefficient of the frequency change between all pairs of two evolved replicates was calculated to describe the parallelism of the evolution at these loci. An average across loci is used to obtain a general parallelism. 100 traits with 10 evolution replicates have been performed for each scenario. With five contributing loci, the genomic evolution of the contributing loci is more parallel across the 10 evolved replicates compared to the case when with 100 contributing loci.

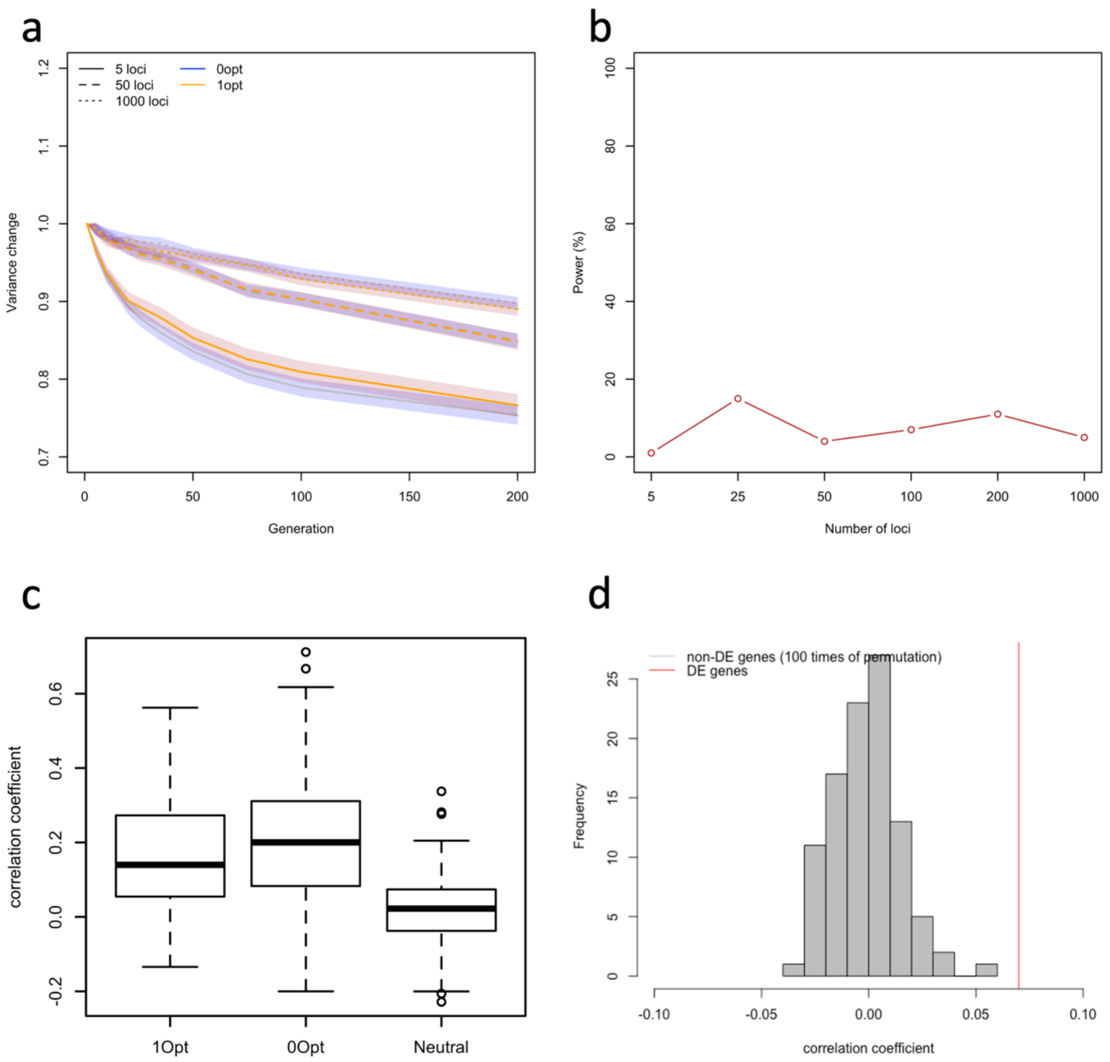


**Supplementary Figure S13. Stabilizing selection without shift in trait optimum. a.** Changes in phenotypic variance (y-axis) across 200 generations (x-axis) of evolution with stabilizing selection either with a shift in trait optimum ($1\sigma^2_{anc}$; orange) or without ($0\sigma^2_{anc}$; blue). The average change in variance of 1000 traits (F) is calculated as the ratio of the phenotypic variance between each evolved time point (generation x) and the ancestral state. The translucent band indicates the 95% confidence interval for 1000 traits. The simulated traits are controlled by a different number of loci (5 loci, 50 loci and 1000 loci). **b.** Power to detect a significantly different variance change between a group of traits with ($1\sigma^2_{anc}$) and without ($0\sigma^2_{anc}$) shift in trait optimum for a different number of loci (x-axis). **c.** Correlation of variance changes in two populations evolved under stabilizing selection with shift in trait optimum ($1\sigma^2_{anc}$), without shift in trait optimum ($0\sigma^2_{anc}$) and under neutrality. The correlation under stabilizing selection is very similar independent of whether a shift in trait optimum is assumed (p-value > 0.05), but much higher than the correlation under neutrality (p-value < 0.05). **d.**

Observed correlation of variance change between two evolved replicates for DE and non-DE genes in the empirical data. Pearson's correlation coefficient of variance changes (log(F)) for DE genes across two evolution replicates was calculated ($r=0.07$; red line). The correlation coefficient is significantly higher than the variance changes for the non-DE genes when we down sampled the non-DE genes to the number of DE genes ($n=4,323$) (grey bar; 100 permutations) ($p$-value $< 0.01$).

**Titles and legend for supplementary files**

**Supplementary file 1. Library information of the sample in this study.** This file provides a list of all sequenced samples and the library information.
